# Graphene-polymer Nanofibers Enable Optically Induced Electrical Maturation in Stem Cell-Derived Cardiomyocytes and Brain Organoids

**DOI:** 10.1101/2024.12.09.627640

**Authors:** Erin LaMontagne, Gisselle Gonzalez, Ritwik Vatsyayan, Blanca Martin-Burgos, Francesca Puppo, Diogo Biagi, Fabio Papes, Shadi A. Dayeh, Alysson R. Muotri, Adam J. Engler

## Abstract

Human pluripotent stem cell (hPSC)-derived electrically excitable cells provide a unique window into development, but they remain electrically immature partially due to the lack of chronic stimulation. Here, we fabricated electrospun polymer nanofibers containing light-reactive reduced graphene oxide (rGO) as part of a new classes of on-demand, electrically active biomaterials to enhance cell function. Fiber size, stiffness, and electrical conductivity varied with rGO concentration, which impacted hPSC-derived cardiomyocyte and neuron responses; with acute light stimulation, cardiomyocytes exhibited increased, synchronous calcium handling, and neurons showed more calcium peaks with higher frequency. Chronic, repetitive nanofiber light stimulation caused brain organoids to become increasingly electrically active and to activate photoreceptor pathways. This work outlines a tunable method where electrical cell functions can be titrated with rGO fibers and light stimulation, and it suggests that repetitive light stimulation may provide a novel method for retinal differentiation.

**HIGHLIGHTS:**

- Electrospun graphene-polymer nanofibers electrically respond to light stimulation
- Light reactive graphene nanofibers stimulate electrically excitable cells in real-time
- Stem cell-derived cardiomyocytes and neurons on nanofibers functionally improve
- Light-training of brain organoids induces retinal and excitable neuron maturation

## 1. Introduction

Human pluripotent stem cell (hPSC)-derived cells have emerged as a promising avenue for modeling development, understanding diseases, and advancing medical therapies. However, conventional differentiation methods using small molecules to guide genetic expression yield functionally immature cells. This is particularly evident for hPSC-derived electrically excitable cells (EECs), which frequently exhibit suboptimal electrophysiology characterized by weak action potentials,^1,2^ low firing rates,^3–5^ naïve network oscillations,^6^ and asynchronous signals^3,4^ compared to EECs *in vivo*.

Conductive biomaterials can propagate electric signals through the microenvironment, and when those biomaterials act as scaffolds for hPSC-derived EECs, they can stimulate EECs, improving their functional matrurity.^7,8^ Among these materials, reduced graphene oxide (rGO) has unique optoelectric properties that can induce electrical action potentials.^8^ rGO is composed of a single sheet of carbon atoms that is very thermally (up to 2,600 W/m-K) and electrically conductive (up to 6,300 S/m).^9–11^ It can also convert light into an electric signal within femtoseconds due to its hot charge carrier properties that facilitate energy transitions and photodetection.^12^ These favorable properties position rGO as an ideal candidate as a scaffold for hPSC-derived EECs. However, rGO’s two-dimensional (2D) structure make it difficult to incorporate into cellular cultures, particularly three-dimensional (3D) spheroid cultures. rGO also fragments into flakes ranging in size from nanometers to millimeters,^13^ and can exhibit a range of hydrophobic properties compared to GO or pristine graphene.^14,15^ To engineer rGO to be more suitable to cell culture, rGO flake-polymer mixtures can be electrospun into nanofibers.^15^ These rGO-polymer nanofibers retain conductivity from rGO^15^ while immobilizing and dispersing rGO flakes in 3D space, enabling rGO’s application in 3D microenvironments. However, the degree to which titratable rGO-polymer material properties influence hPSC maturation and function into EECs is not clear. To address the critical need for supporting functionally mature cultures and to exploit the favorable properties of rGO, we fabricated and implemented conductive, light-activated rGO-poly(viny) alcohol (PVA) nanofibers to electrically stimulate hPSC-derived cardiomyocytes and neurons and brain organoids.

We performed a systematic study to evaluate the effect of rGO concentration on rGO-PVA fiber size, stiffness, and conductive properties. We then demonstrated that 2D EECs cultured on these fibers respond to acute light stimulation with increased and synchronous calcium activity in cardiomyocytes and increased number of calcium spikes and greater neuron spike frequency. Importantly, we performed repetitive, daily “light-training” on brain organoids with rGO-PVA nanofibers and show that the organoids become more electrically responsive over time, with greater bursting activity across more neurons. Through transcriptomic analysis and immunofluorescent staining of these organoids, we discover that repetitive light training induces photoreceptor cell differentiation and excitable neuron maturation in brain organoids with rGO-PVA nanofibers. Collectively, these results demonstrate that acute light stimulation of hPSC-derived EECs with rGO-PVA nanofibers can produce on-demand electrical responses and that long-term light training can enhance electrical activity and promote the emergence of specific cell populations in 3D brain organoids.

## 2. Materials and Methods

### 2.1 PVA-rGO electrospun scaffold fabrication

A solution of 20% (w/v) PVA (Sigma-Aldrich, 363146) and 0.25% (w/v) Triton X-100 (Sigma-Aldrich, X100) in deionized (DI) water was mixed for 3 hours via magnetic stirring at 130 °C. PVA-rGO nanocomposites were made by diluting 20% (w/v) PVA solutions with rGO flakes (Nanotools Bioscience) suspended in DI water, to a final concentration of 10% (w/v) PVA with the desired concentration of rGO. Nanocomposite solutions were vortexed for 1 hour and sonicated prior to electrospinning, up to 4 hours. 18 mm (VWR, 48382-04) and 5 mm (Electron Microscopy Sciences, 7229605) glass coverslips were prepared by plasma etching with oxygen for 1 minute (GLOW Plasma System, Glow Research) to promote even fiber collection on glass surface. Nanocomposite solutions were electrospun onto coverslips using an Inovenso Nanospinner (KTP700 Basic) electrospinner with the following parameters: 0.8 mm nozzle, 0.5mL total solution volume for 18mm coverslips and 0.2mL total solution volume for 5mm coverslips, 20 cm working distance, 30% humidity, 32-35 kV, and 0.4-0.8 mL/hr flow rate (higher voltage and lower flow rates used to electrospin higher rGO concentrations). Scaffolds were crosslinked in a glass chamber (Chemglass, AF-0556) containing 0.2M glutaraldehyde (ThermoFisher Scientific, O2957-1) for 30 minutes at 80°C. Then, scaffolds were removed from the chamber and washed three times in 1% (w/v) glycine (Sigma-Aldrich, G7126) solution for 30 minutes each to inactivate unreacted glutaraldehyde and subsequently three times in DI water for 10 minutes each.^73^

### 2.2 PVA-rGO electrospun nanofiber fabrication

3-4 crosslinked 18mm scaffolds were submerged in water and scraped from the glass coverslips using a pipette tip. The scaffolds were then frozen in DI water in cryomolds at -80 °C overnight. Frozen scaffolds were sectioned on a cryostat (Leica CM3050s) at 20 µm. Frozen shavings were collected in a conical tube, filtered through a 100 µm strainer (Fisher Scientific, 22-363-549), re-frozen -80 °C overnight, and lyophilized in a FreeZone freeze dryer (Labconco, 700201000) for 2-3 days, until all water was removed. Individualized nanofibers were then resuspended at 5 mg/mL in D-PBS (Dulbecco’s Modified Eagle Media; Gibco, 11966025).

### 2.3 Fiber sterilization and coating with protein

5mm fibrous scaffolds on glass coverslips were sterilized for 1 hour by UV in a biohood (Nuaire, NU-427) on each side and placed into a 96-well glass bottom plate (Greiner Bio-One, 655892) prior to a final hour of UV sterilization. The scaffolds were then coated with Matrigel (Corning, CLS354234) at a 1:20 dilution and incubated at 37 °C overnight. Similarly, resuspended nanofibers were exposed to 50 mW/cm^2^ of 365 nm UV light for 15 minutes on a TFM-20V High Performance UV Transilluinator (UVP, 49-95-0423-02) and Matrigel was added to the solution at a 1:100 dilution, mixed by inversion, and incubated at 37 °C overnight.

### 2.4 Fiber imaging and measurements

18 mm fibrous scaffolds were coated sputter coated with iridium by a Emitech Sputter Coater (K575X) for 8 seconds at 85 mA to reduce electron charging and imaged via scanning electron microscopy (SEM) on a Zeiss Sigma 500 scanning electron microscope (SEM) with the following parameters: 3 kV accelerating voltage, 30 µm objective aperture, and 5,000 and 15,000 X magnifications. ImageJ was used to measure nanofiber diameters from SEM images, excluding merged fibers. Suspended nanofibers were imaged on glass slides via light microscopy on a Zeiss Axio Observer and nanofiber lengths were measured using the contour tool in ImageJ along the full length of the fiber curves. Non-individualized nanofiber clusters were measured across their longest lengths.

### 2.5 Bulk stiffness measurements

18 mm fibrous scaffolds were soaked in D-PBS for 1 hour and measured on an Oxford Instruments MFP-3D-Bio atomic force microscope (AFM). A cantilever with a 45 µm size, 0.03 N/m probe (Novascan) and the following parameters were used to generate force maps: 3 µm force distance, 2 µm/s velocity, 0.33 Hz scale rate, 8 Hz sampling rate, and 1 nN trigger point. 20×20 µm force maps with 20 points per map were analyzed by fitting a Hertz model to the curves.^74^

### 2.6 Conductivity measurements

Conductivity measurements were taken of fibrous scaffolds (0.5 mL total electrospun volume) deposited and crosslinked on silicon (Si) wafers (Boron doped p-type Si wafers, 500 µm thick, 10 S/m electrical conductivity). Si wafers were cut with a Disco Automatic Dicing Saw 3220 into 1×1 cm squares and then cleaned using RCA-1 (which is a solution of H_2_O:H_2_O_2_:NH_4_OH in a 5:1:1 ratio by volume) and RCA-2 (which is a solution of H_2_O:H_2_O_2_:HCl in a 5:1:1 ratio by volume), followed by dipping into HF, to remove organic and ionic contaminants and surface oxides from the surface of the wafer. After depositing and preparing the fibers, 4-point resistivity measurements were performed using a Six-Point-Probe-Meter Model 101C (Four Dimensions USA) to produce sheet resistance measurements. Control measurements on PVA fibers were first performed showing insulating characteristics. Therefore, the Si wafer was essentially insulated from the scaffolds. Light activation was supplied during relevant measurements using a PhotonMaker (Nanotools Bioscience) with the following parameters: 452 nm wavelength light; 100% intensity. Sheet thickness was measured by cutting the scaffolds and measuring the height of the scaffold using a Dektak XT Stylus Profilometer (Bruker) with the following parameters: 12.5 µm radius stylus tip, 3 mg stylus force, and 6.5 µm range. Material conductivity was calculated as the inverse of the product of the sheet resistance and sheet thickness.

### 2.7 hPSC-derived cardiomyocyte differentiation and cell seeding

RUES2 hPSCs were cultured on Matrigel-coated 6-well plates, fed daily with mTeSR 1 (Stemcell Technologies, 85850), and chemically passaged using Versene (Gibco, 15040066). The cardiomyocyte differentiation was conducted using the GiWi cardiac differentiation protocol.^25^ At day 50, cardiomyocytes were dissociated by incubation in 4 mg/mL collagenase IV (Gibco, 17104-019) for 50 minutes at 37 °C followed by incubation in 0.25% Trypsin-EDTA (Gibco, 25200) for 25 minutes at 37 °C. The dissociation reagents were deactivated with RPMI (Gibco, 11875-903) + B27 with insulin (ThermoFisher Scientific, 11875-093) + 20% (v/v) FBS (Omega Scientific, FB-01) + 5 µM Y-27632 dihydrochloride (ROCKi; LC Laboratories, Y-5301) and cells were centrifuged at 400 xg for 10 minutes. Cardiomyocytes were resuspended in the same medium and seeded on sterilized, Matrigel-coated 5 mm scaffolds in a 96-well glass bottom plate at 32,000 cells per well, aiming at the center of the well, enough to form a monolayer over the scaffold. The cells remained undisturbed for 2 days, and then were fed every other day with RPMI + B27 with insulin until assayed.

### 2.8 hPSC-derived brain organoid differentiation

H9 hPSCs were cultured on Matrigel-coated 6-well plates, fed every other day with mTeSR Plus (Stemcell Technologies, 100-0276), and chemically passaged using ReLeSR (Stemcell Technologies, 100-0483). The brain organoid differentiation was modified from the cortical organoid protocol described in Trujillo et al^6^ and Fitzgerald et al.^75^ A 24-well AggreWell 800 plate (Stemcell Technologies, 34811) was prepared by adding Anti-Adherence Rinsing Solution (Stemcell Technologies, 07010) to each well, centrifuging the plate at 1300 xg for 5 minutes, and rinsing each well with D-PBS. On Day 1, hPSCs were dissociated into single cells Accutase (Stemcell Technologies, 07920) in D-PBS at a 1:1 ratio for 10 minutes at 37 °C. The cells were then centrifuged for 3 minutes at 150 x *g* and the pellet was resuspended in mTeSR Plus supplemented + 10 μM SB431542 (SB; Stemgent, 04-0010-base) + 1 μM Dorsomorphin (Dorso; R&D Systems, 3093) + 5 μM ROCKi). 2.5×10^6^ cells were transferred to each well of the prepared AggreWell 800 plate, and the plate was centrifuged at 1300 xg for 5 minutes. The AggreWell plate was incubated at 37 °C overnight. On day 2, embryonic bodies were transferred from each used well of the AggreWell plate to each well of a 6-well plate in fresh mTeSR Plus + 10 μM SB + 1 μM Dorso and kept in suspension on an orbital shaker (90 rpm rotation) for the duration of the brain organoid differentiation. Media was changed daily through day 6. From days 7-11, media was changed every other day with M1 media [Neurobasal Medium (Life Technologies, 21103049) + 1% GlutaMAX (ThermoFisher Scientific, 35050061) + 1% Gem21 NeuroPlex (Gemini Bio, 400-160-010) + 1% N2 NeuroPlex (Gemini Bio, 400-163-005) + 1% NEAA (ThermoFisher Scientific, 11140050) + 1% PS (ThermoFisher Scientific, 15140122) + 10 μM SB + 1 μM Dorso] and embryoid bodies were split between new wells approximately once during this period to prevent overcrowding. From days 12-18, media was changed every day with M2 media [Neurobasal Medium + 1% GlutaMAX + 1% Gem21 NeuroPlex + 1% NEAA + 1% PS] + 20 ng/mL FGF2 (Peprotech, 100-25). From days 19-25, media was changed every other day with M2 + 20 ng/mL FGF2 + 20 ng/mL EGF (PeproTech, AF-100-15). From days 26-29, half media changes were performed with M3 media [M2 medium + 10 ng/mL BDNF (PeproTech, 450-02) + 10 ng/mL GDNF (PeproTech, 450-10) + 10 ng/mL NT-3 (PeproTech, 450-03) + 200 μM L-ascorbic acid (Sigma-Aldrich, A4403) + 1 mM dibutyryl-cAMP (Stemcell Technologies, 100-0244)] + 20 ng/mL FGF2. From days 30-35, media was changed every 3-4 days with M3 media. From days 36-42, media was changed every 3-4 days with M2.5 media [M3 medium with half the concertation of factors]. From day 43 on, brain organoids were maintained in M2 media, media changes every 3-4 days.

### 2.9 Neuron isolation from dissociated brain organoids and cell seeding

2-month-old brain organoids were first moved to a new 6-well plate. Old media was removed, and the organoids were washed with 0.5M EDTA (Invitrogen, 15575020) in D-PBS, then incubated in TrypLE Select Enzyme 10X (ThermoFisher Scientific, A1217701) at 37 °C on an orbital shaker (90 rpm) for 70 minutes. The dissociation reaction was quenched with a blocking solution of 2 μM CaCl_2_ (Sigma-Aldrich, 21115) + 10 μM ROCKi + 1% BSA (Sigma-Aldrich, A9576) + 400 U/mL DNAse (Worthington Biomedical, LK003172) in M2 media and brain organoids were mechanically dissociated by gently pipetting up and down approximately 15-50 times. Cells were strained with a 70 μm strainer (FisherScientific, 22-363-548), centrifuged at 300 xg for 5 minutes, and resuspended in M2 media + 5 μM ROCKi + 5% FBS. Cells were seeded on sterilized, Matrigel-coated 5 mm scaffolds in a 96-well glass bottom plate at 150,000 cells per well, aiming at the center of the well, enough to form a monolayer over the scaffold. The cells remained undisturbed for 2 days, and then were fed every other day with M2. The day before assaying, they were fed with BrainPhys Neuronal Medium (Stemcell Technologies, 05790) + 1% Gem21 NeuroPlex.

### 2.10 Calcium imaging and acute light stimulation

Calcium imaging was performed on hPSC-derived cardiomyocytes and neurons from dissociated brain organoids seeded on 5 mm fibrous scaffolds with varying rGO concentrations. After 2 weeks of being cultured on scaffolds, cells were incubated with their respective culture media + 2 µM fluo-4 AM ester (Biotium, 50018) for 15 minutes at 37 °C protected from light. 30 second recordings were captured using a Zeiss Axio Observer with a 20x objective. Acute light stimulation was supplied 1-3 times during relevant recordings using a PhotonMaker (Nanotools Bioscience) with the following parameters: 452 nm wavelength light; 100% intensity; 2 consecutive cycles of 12 repetitions; 3 Hz pulse frequency; 25 ms pulse duration. Stimulation was recorded by live video microscopy for 30 sec durations. Analysis was performed over the entire movie duration by first using the contour tool in ImageJ to outline individual cells (only the soma for neurons) and to generate a kymograph of the fluorescence flux over the recording duration. Calcium imaging measurements were analyzed using a custom R script, available on Github (https://github.com/eelamonta/Graphene-Project). Peak amplitude was calculated as the change in fluorescence intensity from local min to local max, and time-to-peak was determined as the time between local min and local max. Data parameters and replicates are annotated in Tables S1 and S2.

### 2.11 MEA seeding and fiber-coating

CytoView MEA 48 plates (Axion Biosystems, M768-tMEA-48B) were first sterilized by incubating the wells with 1% Terg-a-zyme (Sigma-Aldrich, 7273287) in sterile DI water for 2 hours. Terg-a-zyme was removed from the wells and the wells rinsed with sterile DI water. The wells were sterilized a second time by incubating 70% ethanol (Sigma-Aldrich, E7023) in sterile DI water for 30 minutes. Ethanol was removed from the wells and the wells rinsed with sterile DI water. M2 media was added to the wells and incubated for at least 48 hours at 37 °C to pre-condition the surface. After pre-conditioning, wells were rinsed with sterile DI water, air-dried, and a solution 0.07% PEI (Sigma-Aldrich, P3143) in 1X borate buffer (ThermoFisher Scientific, 28341) sterilized through a 0.22 μm filter (Millipore Sigma, SCGP00525) was added to the wells and incubated at 37 °C for 1 hour. Then the wells were rinsed with sterile DI water, air-dried, and coated with 0.4 mg/mL laminin (Life Technologies, 23017015) in D-PBS by pipetting a small volume directly onto the electrodes, without allowing the solution to spread to the edges of the well. The plates were then incubated at 37 °C for 1 hour. M2 media was added to the wells, followed by 2-4 organoids between 60-90 days of differentiation and then 100 μg of sterilized, Matrigel-coated nanofibers were added to each well. Micropipette tips were used the move the organoids onto the electrodes, taking care not to scratch the wells’ surfaces. The plates were then incubated at 37 °C for 5 days to allow organoids to attach. Half media changes were performed every 4-5 days, taking care not to disturb the brain organoids as networks were forming.

### 2.12 MEA light stimulation and recording

MEA recordings were performed on the Maestro Pro (Axion Biosystems) approximately 2 weeks after seeding the brain organoids on MEA plates, every 4-5 days (the day after media change) for 8 weeks. 1 hour of daily light stimulation was produced using the Lumos (Axion Biosystems) with the following parameters: both blue and green lights at 100% intensity (>3.9 mW/mm^2^ and >1.9 mW/mm^2^, respectively, per manufacturer), 1-2 Hz pulse frequency, 1 ms pulse rise, 10 ms pulse duration, 5 ms pulse off. Light frequency was increased from 1 to 2 Hz in 0.3 Hz intervals over the 8 weeks of light training. 5-minute recordings were made before light stimulation, during the last 5 minutes of light simulation, and 5 minutes directly after light stimulation for the light-trained MEA plates. Sustained, long-term light responses were the pre-light recordings. Acute, short-term light responses were directly after the hour of light training. Control plates were never exposed to light stimulation and a single 5-minute recording was made on recording day for these plates.

### 2.13 RNA isolation, sequencing, and analysis

Brain organoids were scraped from MEA plates using a micropipette tip, moved to conical tubes, rinsed with D-PBS, and pelleted by gravity for 3 minutes. The supernatant was aspirated and RNA was extracted using the RNeasy Mini Kit (Qiagen, 74104). RNA quality was measured using an Agilent TapeStation 4200 and only RNA with RINe values greater than 7.0 were used for sequencing. Illumina Stranded mRNA bulk RNA sequencing libraries were sequenced on an Illumina NovaSeq 6000 with paired end reads (PE150). Sequencing quality was assessed using FASTQC^76^ and adapters were trimmed using Trim Galore!.^77^ Reads were transcriptionally mapped to the human reference genome hg38 using kallisto^78^ and feature aggregation was preformed using tximport^79^. Batch correction and normalization of data using log counts per million was achieved using ComBat^80^ via the SVA package. pheatmap^81^ was used to generate a heatmap from a t-test of normalized gene expression. Differential gene expression between the assigned groups was determined with limma-voom^82,83^ and plots were generated using ggVennDiagram^84,85^ and ggplot2^86^ with the following cutoffs: absolute value of log fold change greater than 0.6, p-value less than 0.05, and absolute value of normalized log expression greater than 3. Of note, the Banjamini-Hochberg adjusted p-values for the majority of DEGs were above 0.05, hence we analyzed the sequencing results using the unadjusted p-values as indicators of significance. EnrichGO and gseGO via clusterProfiler^87,88^ were used for pathway analysis of differentially expressed genes. RNAseq analysis was accomplished using a custom R script, available on Github (https://github.com/eelamonta/Graphene-Project).

### 2.14 Brain organoid immunofluorescence staining and imaging

Organoids were rinsed with D-PBS and fixed in 4% paraformaldehyde (ThermoFisher Scientific, J19943K2) at 4 °C for 4 hours. Brain organoids were then incubated in 30% sucrose (Sigma-Aldrich, S0389) in D-PBS overnight. Brain organoids were then mounted in OCT (Sakura Finetek USA, 4583) and frozen at -20 °C overnight before being cryosectioned at 20 μm. Slides were stored at -20 °C until staining. In preparation for staining, slides were dried for 10-20 minutes, rinsed in D-PBS to remove the OCT medium, and blocked in a solution of 2% BSA and 0.1% Triton X-100 in D-PBS for 2-3 hours. Slides were then incubated in primary antibodies diluted in blocking solution at 4 °C overnight. After rinsing the slides in D-PBS they were incubated with secondary antibodies diluted in blocking solution for 2-3 hours in the dark, rinsed again in D-PBS, and then counterstained with 1 μg/ml DAPI (Chemometec, 910-3003) in 0.1% Triton X-100 in D-PBS for 45 minutes in the dark. After a final rinse in D-PBS, coverslips were mounted on the slides using ProLong Gold (ThermoFisher Scientific, P36930). Primary antibodies used were: FOXG1 (Millipore, MABD79; 1:500), DCX (Abcam, ab18723; 1:200), MAP2 (Abcam, ab5392; 1:1000), ZO-1 (Invitrogen, 33-9100; 1:100), S-OPSIN (Invitrogen, OSR00219W; 1:500), RCVRN (Millipore, AB5585; 1:2000), CRX (RD Systems, AF7085; 1:100), RHO (Abcam, ab5417; 1:200), VGLUT1 (Synaptic Systems, 135311; 1:100), and GAD65+67 (Abcam, ab11070; 1:200). Secondary antibodies (Invitrogen; 1:500) used were: Goat anti-Rabbit Alexa Flour 488 (A11034), Donkey anti-Mouse Alex Flour 555 (A31553), Goat anti-Rabbit Alexa Flour 647 (A21244), Donkey anti-Mouse Alexa Flour (A21202), Donkey anti-Mouse Alexa Flour 647 (A31571), Donkey anti-Sheep Alexa Fluor 647 (A21448), and Goat anti-Chicken Alexa Flour 647 (A21449).

Slides were imaged on the Dragonfly 600 (Andor Technology) and analyzed using Imaris Image Analysis Software v10.1 (Oxford Instruments). Vesicle proteins were quantified using Imaris by quantifying spheres of a specific seed size (3.5 μm for DAPI+ nuclei and 0.2 μm for VGLUT+ and GAD65+67+ puncta) and voxels of a specific threshold (15.5 for DAPI+ nuclei and 25 for VGLUT+ and GAD65+67+ puncta). Vesicle protein quantities were then normalized to the number of DAPI+ nuclei.

### 2.15 Statistical analysis

Statistical test and sample size (n) are reported in figure descriptions. Statistical analyses were performed using R v4.3.3 in R studio v2023.12.1+402 with the ggstats^89^ and ggpubr^90^ packages. Box plots created via ggplot2^86^ show the median values as a line, the 1^st^ to 3^rd^ interquartile range as the box, the maxima and minima (defined as values within 1.5 times the interquartile range) as whiskers, and outliers as points. Statistical analyses were made using Kruskal-Wallis and Wilcoxon Tests. In plots comparing pre-light to light stimulation or post-light values, statistical comparisons were only made between each pre-light condition, each light stimulation or post-light condition, and rGO concentration-matched conditions. Significance was defined as * p<0.5, ** p<0.01, *** p<0.001, and **** p<0.0001. If significance between any of the compared groups is not graphically described in a figure that was statistically analyzed, the comparison was not significant.

## 3. Results

### 3.1 Electrospun rGO-PVA fiber diameter, stiffness, and conductivity scales with rGO concentration and their conductivity increases with acute light stimulation

Fibrous scaffolds were generated by electrospinning and crosslinking nanocomposites of PVA with varying concentrations of rGO flakes (Fig. 1a). Scaffolds were primarily composed of fibers between 250-500 nm in diameter, about the same diameter as neurite process^9^, with fiber diameter decreasing with increasing rGO concentration (Fig. 1b), potentially a result of changing solution viscosity during electrospinning.^15,16^ Notably, 0.1% rGO fibers were larger in diameter than 0.01% rGO fibers, and while parameters changed to enable eletrospinning at all rGO concentrations, substantial changes may have been necessary at higher concentrations due to greater rGO flake coagulation. The bulk fiber modulus of the scaffolds was also evaluated, since material stiffness is known to influence PSC differentiation^17^, cardiomyocyte structure and electrophysiology,^18^ and neural network formation.^19,20^ Scaffold Young’s Modulus scaled with rGO concentration in the range of stiff yet compliant tissues (Fig. 1c). Although this is substantially greater than the stiffness of *in vivo* tissues, studies have shown that neurons grown in artificial environments are capable of forming neurite outgrowths on ultra-stiff materials including graphene^20,21^. Cardiomyocytes, however, prefer substrate stiffness within a normal physiological range^18,22^. Most importantly, electrical conductivity of the scaffolds similarly varied with rGO concentration up to 3×10^-3^ S/m (Fig. 1d) and conductivity of rGO-containing scaffolds increased up to 6-fold when acutely activated with light (Fig. 1e); PVA-only scaffolds were consistently highly insulating, with unstable conductivity values at or below the limit of our measurable range.

**Fig. 1.**
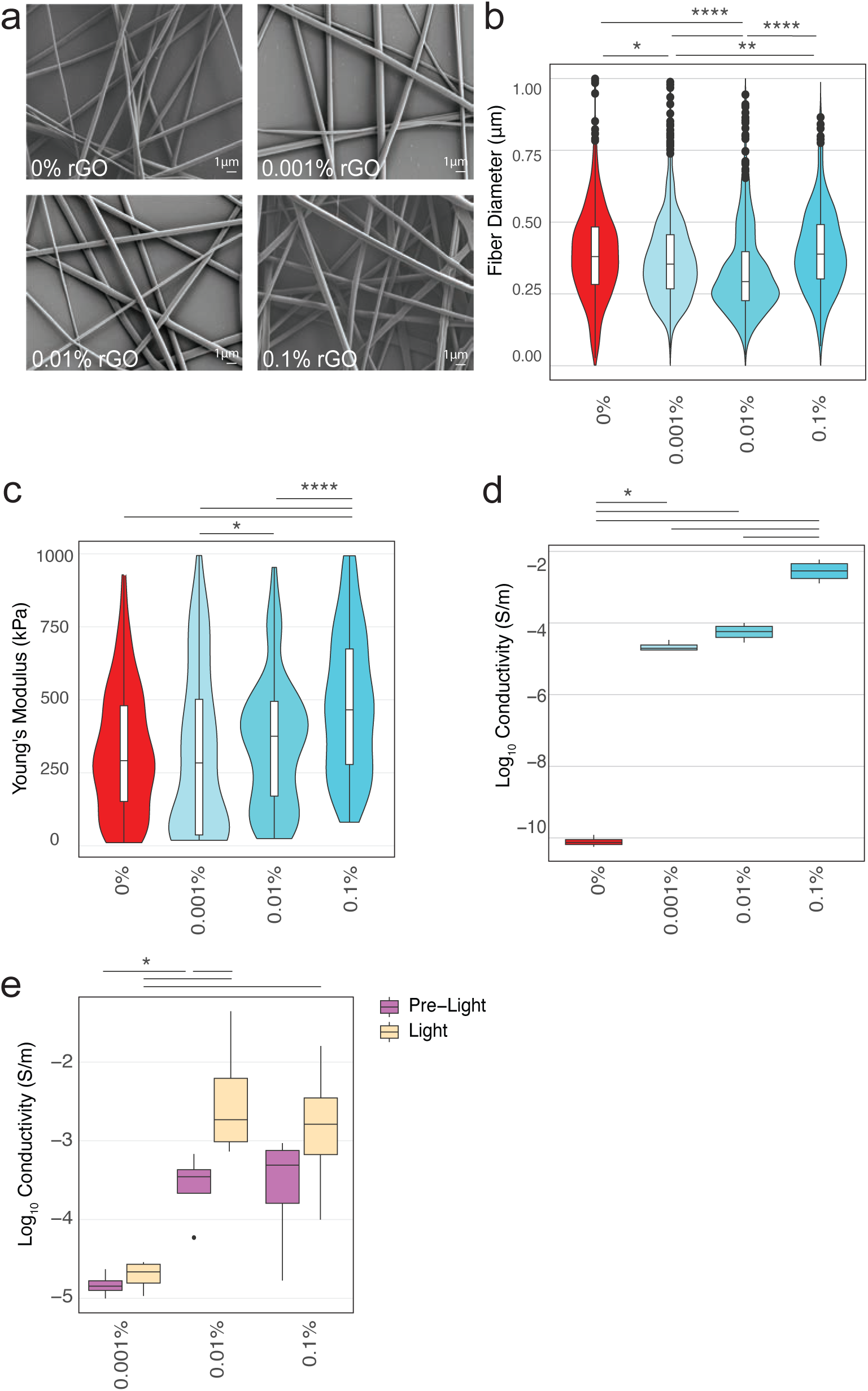
Physical characterization of electrospun fibrous scaffolds. (**a)** SEM images of 10% PVA scaffolds containing 0%, 0.001%, 0.01%, and 0.1% rGO. Magnification = 5,000 X. Scale bar = 1 µm. **(b)** Fiber diameter measurements from SEM images (N = 3-5 scaffolds per condition; n = 25-100 measurements per scaffold). Median fiber diameters of 10% PVA scaffolds containing 0%, 0.001%, 0.01%, and 0.1% rGO were 380, 356, 294, and 391 nm, respectively. Kruskal-Wallis, p<2.2e-16. **(c)** Young’s Moduli of scaffolds (N = 3-4 scaffolds per condition; n = 25-50 measurements per scaffold). Median Young’s Moduli of 10% PVA scaffolds containing 0%, 0.001%, 0.01%, and 0.1% rGO were 291, 283, 375, and 465 kPa, respectively. Kruskal-Wallis, p=8.7e-9. **(d)** Conductivity of scaffolds (N = 4 scaffolds per condition; n = average of 6-8 measurements per scaffold). Median conductivities of 10% PVA scaffolds containing 0%, 0.001%, 0.01%, and 0.1% rGO were 7.49e-11, 1.98e-5, 5.95e-5, and 3.06e-3 S/m, respectively. Kruskal-Wallis, p=0.0032. **(e)** Conductivity of scaffolds before and during light stimulation (N = 4 scaffolds per condition; n = average of 6-8 measurements per scaffold). Median conductivities of 10% PVA scaffolds containing 0.001%, 0.01%, and 0.1% rGO pre-light stimulation were 1.42e-5, 3.5e-4, and 5.22e-4 S/m, respectively; 10% PVA scaffolds containing 0.001%, 0.01%, and 0.1% rGO during stimulation were 2.21e-5, 2.14e-3, and 1.68e-3 S/m, respectively. Kruskal-Wallis, p=0.0036. Statistical analyses were made using Kruskal-Wallis and Wilcoxon Tests and significance is defined as * p<0.5, ** p<0.01, *** p<0.001, and **** p<0.0001.

### 3.2 2D hPSC-derived cardiomyocytes and neurons on rGO-PVA scaffolds show improved calcium handing in response to acute light stimulation

To evaluate the effects of rGO-PVA nanofibers on EEC electrical function, we first analyzed calcium handling in 2D hPSC-derived EECs cultured on fibrous scaffolds prior to and during acute light stimulation. D50 hPSC-derived cardiomyocytes were cultured on nanofiber scaffolds for 2 weeks after differentiation (Fig. 2a) and selectively exposed to light stimulation (Fig. 2b-h). Cardiomyocytes cultured on rGO-PVA scaffolds were morphologically normal, i.e., an elongated phenotype (Fig. 2d). In the absence of light, the number of spikes and the synchronicity between cells, as described by the Spearman Rank Correlation Coefficient, both varied with rGO concentration (Fig. S1); without rGO present in PVA fibers, cells were largely uncoordinated, as describe by the Spearman Rank Correlation Coefficient (Fig. S1c). However, there was a clear response of cardiomyocytes on rGO-PVA scaffolds to acute light stimulation with improved calcium spike synchronicity (Fig. 2g-h), suggesting rapid cellular response to light-activation of rGO. While significantly higher than all other conditions, the Spearman Rank Correlation Coefficient suggests that two independently synchronous cardiomyocyte populations could exist on the same scaffold with light stimulation (Fig. 2g), perhaps resulting from heterogenous distribution of rGO throughout PVA fibers. Greater peak amplitude and faster time-to-peak also indicated heightened electrical activity in light-stimulated cardiomyocytes on PVA scaffolds with increasing rGO (Fig. 2h and Table S1); in the absence of rGO, light was ineffective at producing coordinated calcium spikes (Fig. 2g-h).

**Fig. 2.**
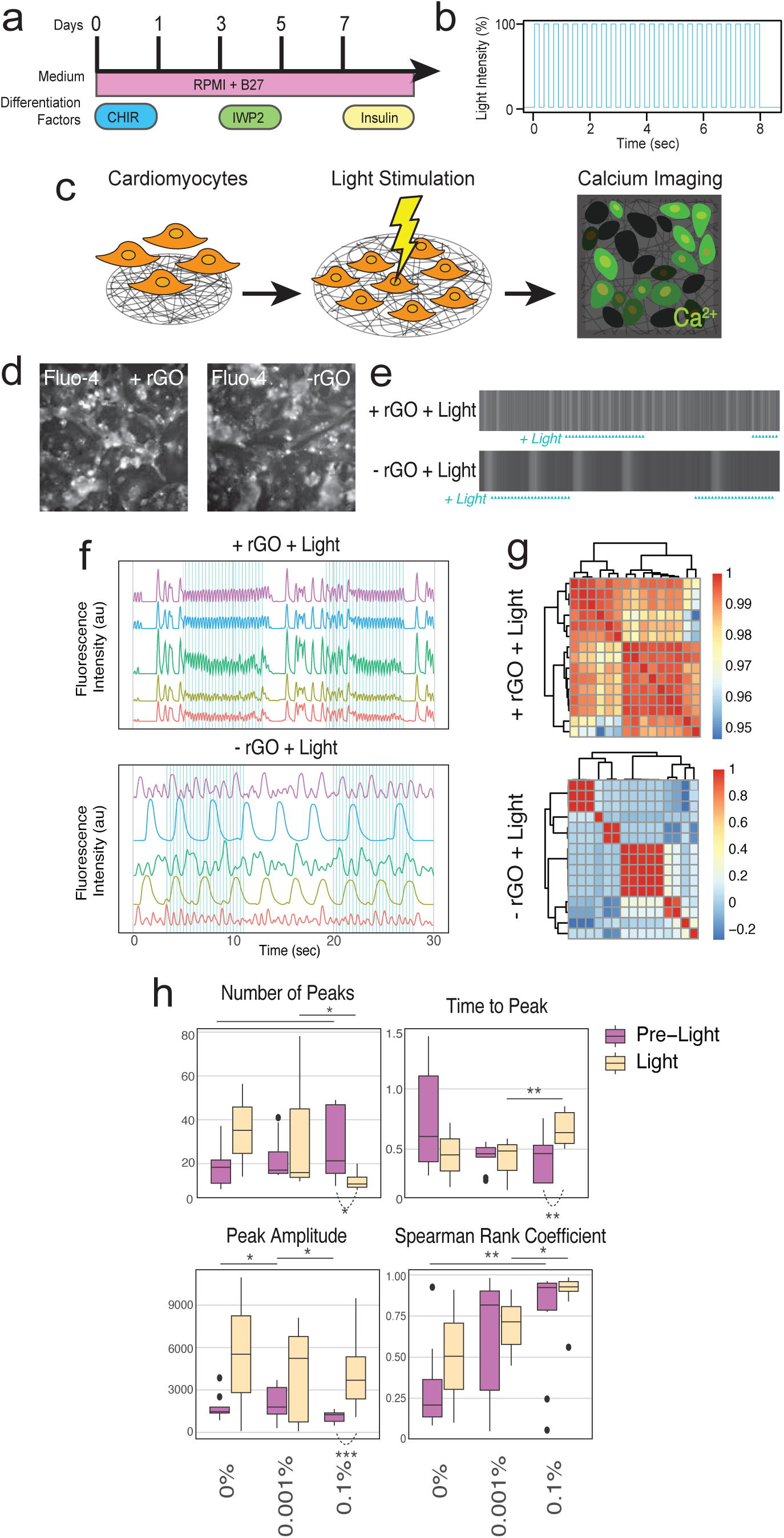
Cardiomyocytes on rGO-PVA scaffolds are more synchronous and exhibit distinct calcium signals in response to acute light stimulation. **(a)** Timeline of PSC-derived cardiomyocyte differentiation. **(b)** Light stimulation profile used to acutely stimulate cells on fibrous scaffolds (452 nm wavelength light; 2 consecutive cycles of 12 repetitions; 3 Hz pulse frequency; 25 ms pulse duration). **(c)** Schematic of cardiomyocyte calcium handling assay. After D50 of differentiation, PSC-derived cardiomyocytes are cultured on scaffolds for 2 weeks prior to calcium imaging with and without acute light stimulation. **(d)** Fluorescent images of cardiomyocytes on fibrous scaffolds during calcium imaging. Fluo-4 is a calcium indicator dye. Elongated cardiomyocytes are outlined in orange. **(e)** Representative kymographs with blue triangles indicating light stimulation time points and changes in brightness indicating calcium flux, (**f)** representative light intensity profiles in arbitrary units to illustrate coordination of spikes and light, and (**g)** heatmaps of Spearman Rank Coefficients of cardiomyocytes on scaffolds with and without rGO and with or without acute light stimulation (light pulses are indicated by carrots in e and vertical blue lines in f). **(h)** Calcium handling metrics of cardiomyocytes on scaffolds containing 0%, 0.001%, and 0.1% rGO with and without acute light stimulation. Time to peak, peak amplitude, and spearman rank coefficients have Kruskal-Wallis p-values of 0.11, 0.047, 0.036, and 0.0056, respectively. Median values are reported in Table S1. (N = 4-5 cardiomyocyte differentiation batches with the number of cells per batch analyzed, n = 10, 9, 12 for 0%, 0.001%, and 0.1% rGO without light, respectively, and n = 2, 7, 8 for 0% 0.001%, and 0.1% rGO with light, respectively). Statistical analyses were made using Kruskal-Wallis and Wilcoxon Tests and significance is defined as * p<0.5, ** p<0.01, *** p<0.001, and **** p<0.0001.

To evaluate if light-activation of rGO-PVA nanofibers can similarly stimulate hPSC-derived neurons, we dissociated D60 brain organoids (Fig. 3a) and cultured the resulting cells on fibrous scaffolds for 2 weeks prior to calcium imaging (Fig. 3b) because studies have shown 3D brain organoids exhibit a higher degree of electrical function than 2D hPSC-derived neurons.^23,24^ Unlike the GiWi differentiation protocol for cardiomyocytes,^25^ brain organoids contain multiple cell types, hence we selected cells that were morphologically typical of neurons, i.e., a small soma with long processes (Fig. 3c) and that produced rapid bursts of calcium spikes, an electrical signal unique to neurons (Fig. 3d and Fig. S2a).

**Fig. 3.**
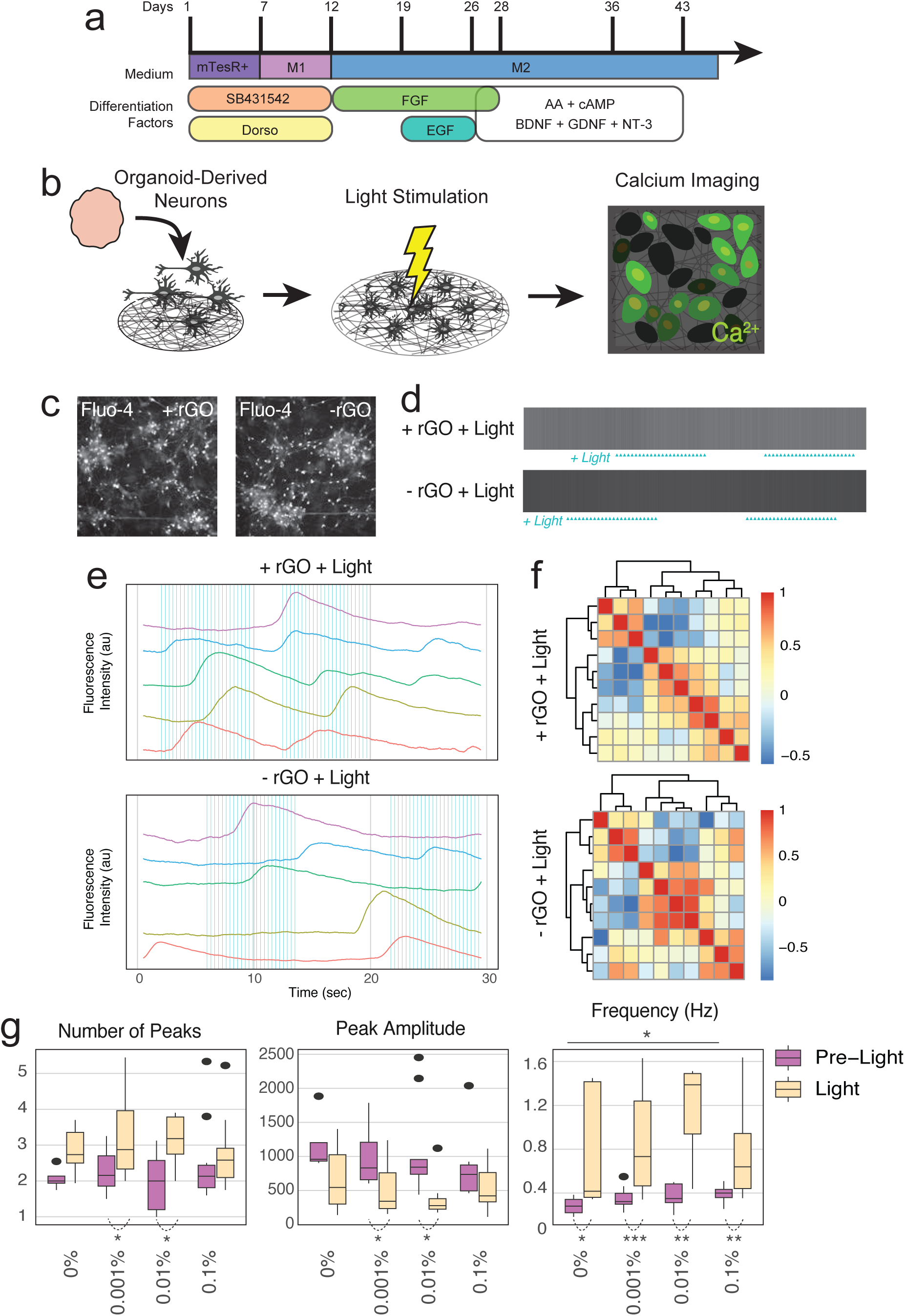
Neurons on rGO-PVA scaffolds generate more calcium spikes in response to acute light stimulation. **(a)** Timeline of PSC-derived brain organoid differentiation. **(b)** Schematic of neuron calcium handling assay. After D60 of brain organoid differentiation, PSC-derived neurons are isolated from dissociated brain organoids and cultured on scaffolds for 2 weeks prior to calcium imaging with and without acute light stimulation. **(c)** Fluorescent images of neurons on fibrous scaffolds during calcium imaging. Fluo-4 is a calcium indicator dye. **(d)** Representative kymographs with blue triangles indicating light stimulation time points and changes in brightness indicating calcium flux, (**e**) representative light intensity profiles in arbitrary units to illustrate coordination of spikes and light, and (**f**) heatmaps of Spearman Rank Coefficients of neurons on scaffolds with and without rGO with or without acute light stimulation (light pulses are indicated by carrots in d and vertical blue lines in e). **(g)** Calcium handling metrics of neurons on scaffolds containing 0%, 0.001%, 0.01%, and 0.1% rGO with and without acute light stimulation. Number of peaks, peak amplitude, and frequency (Hz) have Kruskal-Wallis p-values of 0.023, 0.0042, and 6.9e-6, respectively. Median values are reported in Table S2. (N = 5 brain organoid differentiation batches with the number of cells per batch analyzed, n = 4, 9, 12, 10 for 0% 0.001%, 0.01%, and 0.1% rGO without light, respectively, and n = 6, 11, 7, 12 for 0%,0.001%, 0.01%, and 0.1% rGO with light, respectively). Statistical analyses were made using Kruskal-Wallis and Wilcoxon Tests and significance is defined as * p<0.5, ** p<0.01, *** p<0.001, and **** p<0.0001.

To eliminate potential background noise from light pulses, calcium traces were computationally smoothed, and while erasing fast, singular calcium spikes (Fig. S2b), it enables clean quantification of bursting events, which are visible as “peaks” (Fig. 3e and Fig. S2c). Neurons cultured on scaffolds exhibited calcium events at frequencies around 1 Hz (Fig. 3e, g), and in the absence of light stimulation, no significant difference was evident in calcium handling activity between neurons cultured on scaffolds with versus without rGO (Fig. 3g and Fig. S2c-e). However, more bright puncta were apparent along neuronal processes of cells cultured on rGO-PVA scaffolds, suggesting greater influx of calcium into these cells (Fig. 3c). Upon acute light stimulation, neurons showed significantly improved calcium handling activity denoted by more calcium event peaks and higher frequencies in all rGO-PVA conditions that also scaled with rGO concentration (Fig. 3f-g and Table S2). Unlike cardiomyocytes, neuronal calcium signals were not more synchronous under light stimulation on rGO-PVA scaffolds (Fig. 3f and Fig. S2d). These data together suggest a unique, dose-dependent benefit to the presence of rGO in scaffold fibers, likely due to their structure, their mechanical compliance, and especially their conductive properties.

### 3.3 Light-training 3D brain organoids with rGO-PVA nanofibers improves electrical function over time via photosensitivity

For application into 3D brain organoid culture, scaffolds were further processed by cryo-sectioning into individualized nanofiber suspensions to aide in organoid-fiber incorporation (Fig. S3a). Fiber length was optimized to span 10-80 µm in length (Fig. S3b), which is the length of neurites seen in hPSC-derived neurons *in vitro*.^26^ To facilitate cell adhesion, fibers were coated with Matrigel; rGO-PVA scaffold conductivity generally decreased after Matrigel-coating but conductivity remained significantly higher than PVA-only scaffolds (Fig. S3c). Using these individualized rGO-PVA fibers, we next assessed if light-activation of rGO-PVA nanofibers integrated with brain organoids enhanced electrical maturation. D60 nanofiber-coated brain organoids were seeded on multielectrode arrays (MEAs) and exposed to daily, repetitive light stimulation at incrementally increasing frequencies from 1.33 to 2 Hz over 8 weeks (Fig. 4a-c). After 1-2 weeks of organoid attachment on MEAs, neurite outgrowths protruding from the organoid bodies formed a network across the electrodes and engaged fibers (Fig. 4d). First, we measured fiber-only controls (i.e., in the absence of cells) at various light frequencies to quantify MEA background noise and found that there were constant, low amplitude background “spikes” associated with rGO-PVA nanofiber conditions independent of rGO concentration, the presence of light stimulation, and light stimulation frequency; notably, there were no bursts or network bursts produced by the fiber-only controls (Fig. S4a). Hence, burst and network burst metrics in addition to relative increases in spike counts appear to be reliable indicators of cellular activity by MEA.

**Fig. 4.**
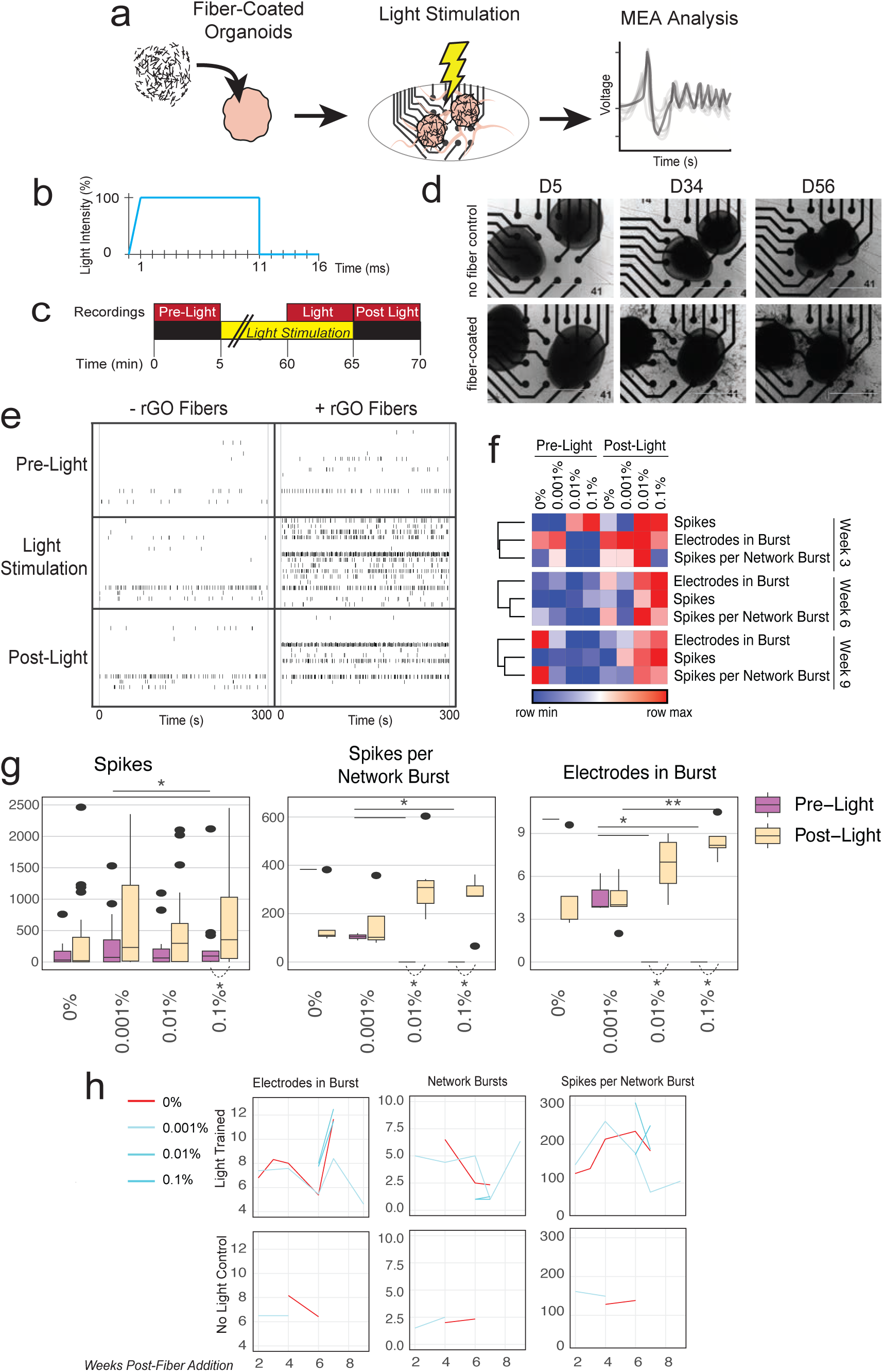
Brain organoids with rGO-PVA nanofibers on MEAs exhibit greater bursting activity between over more electrodes in response to daily acute light stimulation and chronic light training. **(a)** Schematic of brain organoid MEA assay. After D60 of brain organoid differentiation, organoids are coated with nanofibers and cultured on MEA plates where activity is recorded over 8 weeks. **(b)** Light profile used to stimulate brain organoids on MEA (475 nm and 530 nm wavelength light at >3.9 mW/mm^2^ and >1.9 mW/mm^2^, respectively; 1-2 Hz pulse frequency; 1 ms pulse rise, 10 ms pulse duration, 5 ms pulse off). **(c)** MEA activity recording schedule consisting of: a 5 min “pre-light” recording, a 5 min “light stimulation” recording at the end of 1 hr of light stimulation, and a 5 min “post-light” recording taken immediately after light stimulation. **(d)** Brain organoids on MEA showing neurite projections over electrodes 5-, 34-, and 56-days post-organoid seeding. **(e)** Representative raster plots of brain organoids on MEA before, during, and after light stimulation. Each row represents an electrode in an MEA plate well. **(f)** Heatmaps of median values of MEA metrics at 3-, 6-, and 9-weeks post-fiber addition, comparing pre-and post-light recordings. **(g)** Median MEA metrics of brain organoids with nanofibers before and after acute light stimulation over 8 weeks of chronic, daily light training (up to week 9 post-fiber addition). Spikes, spikes per network burst, and electrodes in burst have Kruskal-Wallis p-values of 0.091, 0.0016, and 0.00075, respectively. **(h)** Mean MEA metrics of brain organoids with nanofibers “pre-light” stimulation recordings compared to organoids never exposed to light stimulation over 8 weeks. (N = 3 brain organoid differentiation batches; n = 36-45 MEA wells per condition). MEA metric values are reported in Table S3. Statistical analyses were made using Kruskal-Wallis and Wilcoxon Tests and significance is defined as * p<0.5, ** p<0.01, *** p<0.001, and **** p<0.0001.

Over the course of 8 weeks, MEA signals were recorded before (“pre-light”), at the end of 1 hour of daily light stimulation (“light stimulation”), and after light stimulation ceased (“post-light”) (Fig. 4c). Brain organoids consistently produced the most spikes and activated more electrodes per burst both during and after light stimulation in the presence of rGO-PVA nanofibers, indicating greater neuronal firing activity (Fig. 4e-f). Electrodes that were activated during light stimulation in rGO-PVA conditions remained active after light stimulation subsided (Fig. 4e), suggesting that acute light-induced activation of neurons by rGO evokes sustained electrical activity in brain organoids. MEA recording analysis revealed increased sensitivity to acute light stimulation in brain organoids with rGO-PVA nanofibers over time, denoted by greater numbers of spikes, electrodes in bursts, and spikes per network bursts in post-light recordings at later time points, particularly in with higher (0.01 and 0.1%) rGO conditions (Fig. 4f-g, Fig. S4b, S5, and Tables S3-S4). Repetitive stimulation over time occurs during brain development to build neural circuits,^27,28^ hence we also evaluated the long-lasting impact of chronic light stimulation on neurons, i.e., pre-light recordings of light-trained brain organoids versus unstimulated brain organoids. Light-trained brain organoids exhibited more electrodes involved in bursts, more network bursts, and more spikes per network burst in contrast to unstimulated brain organoids, which rarely produced bursting activity (Fig. 4h and Fig. S6), suggesting that the combination of rGO-PVA fibers and long-term, repetitive light training can significantly improve brain organoid electrical function.

**Fig. 5.**
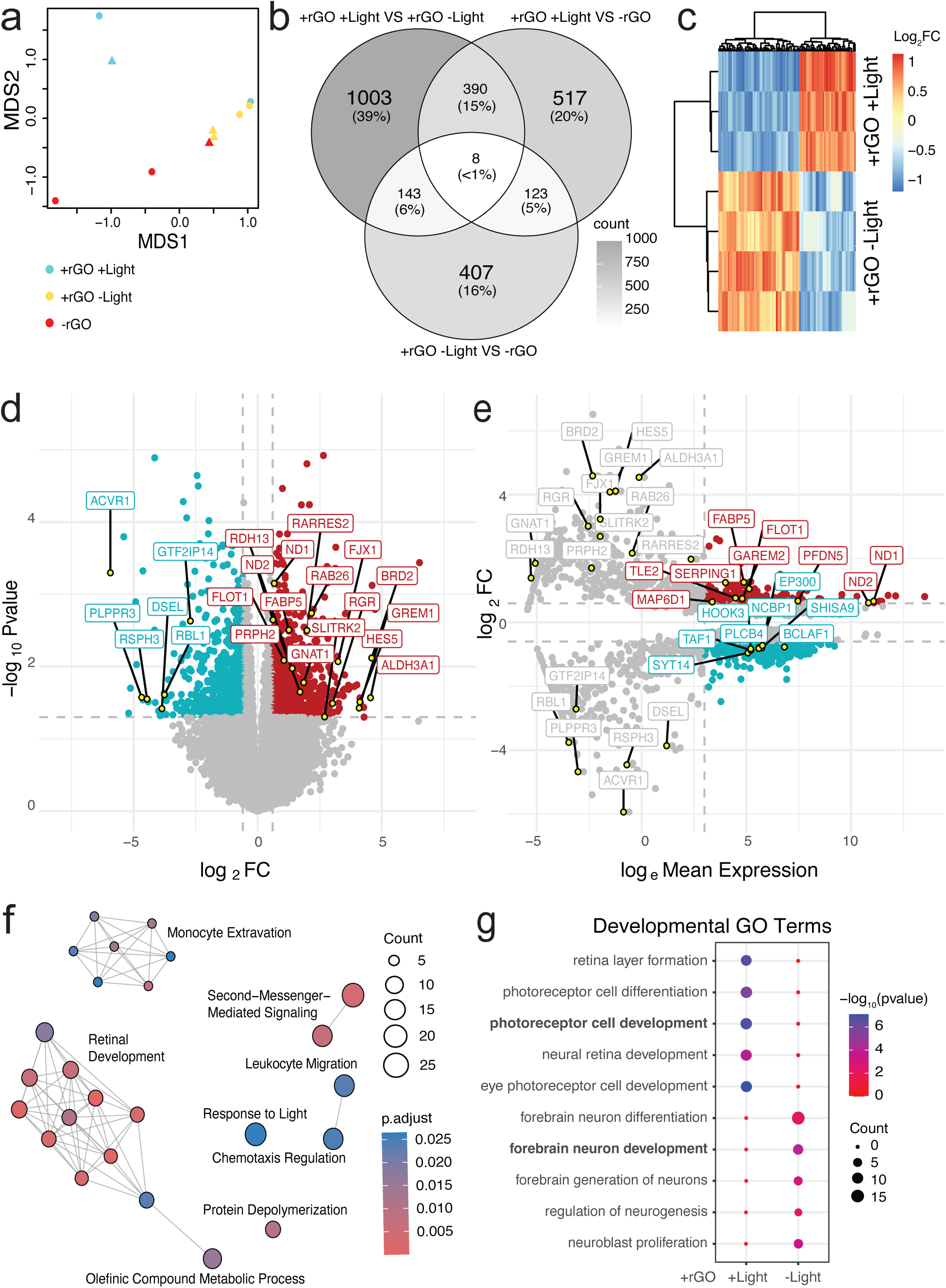
RNA sequencing of light-trained brain organoids with rGO-PVA nanofibers indicate upregulation of retinal development and neuronal maturation pathways. **(a)** MDS plot of RNA transcripts from sequenced brain organoids. Normalized transcript counts are reported in Table S5 (N = 2 brain organoid batches, indicated by triangles and circles; n = 3, 4, 3 for -rGO, +rGO -Light, and +rGO +Light, respectively). **(b)** Venn diagram showing overlapping DEGs between the 3 experimental groups from a. **(c)** Heatmap of the top 100 DEGs between light-trained and unstimulated brain organoids with rGO-PVA nanofibers. Rows represent technical replicates and columns represent individual DEGs. **(d)** Volcano plot of DEGs between light-trained and unstimulated brain organoids with rGO-PVA nanofibers. Cutoffs occur at +/- 0.6 log_2_ fold change and p-values less than 0.05. Labeled DEGs are relevant to brain and eye development. **(e)** MA plot of DEGs and labeled DEGs from d. Cutoffs occur at +/- 0.6 log_2_ fold change and log_e_ mean expression greater than 3. DEGs with p-values less than 0.05 were excluded. **(f)** Emap of top 25 enriched GO terms of DEGs from d. **(g)** Dotplot of enriched GO terms of DEGs from d related to forebrain and retinal development.

### 3.4 Light-training 3D brain organoids with rGO-PVA nanofibers increases expression of excitatory synapse proteins and upregulates retinal differentiation pathways

Improved calcium handling and electrical function in repetitively light-trained, rGO-PVA nanofiber-coated brain organoids should coincide with further maturity at the molecular level. Thus, we performed bulk RNA sequencing on light-trained and unstimulated brain organoids with nanofibers to assess the organoids’ transcriptomes and to identify gene expression changes indicative of specific phenotypes. Excluding ribosomal RNAs, the normalized expression data (Table S5) indicated 3 clusters: light-trained organoids with rGO-PVA nanofibers (+rGO +Light), unstimulated organoids with rGO-PVA nanofibers (+rGO -Light), and organoids with PVA-only nanofibers (-rGO; both light-trained and unstimulated organoids) (Fig 5a). The most differentially expressed genes (DEGs) were found between light-trained and unstimulated brain organoids, both containing rGO-PVA nanofibers (+rGO +Light vs +rGO -Light; 1,003 unique DEGs) (Fig. 5b-c). This suggests that light-activation of rGO caused the largest transcriptomic changes, as opposed to the presence of rGO (407 unique DEGs). DEGs upregulated in light-trained, rGO-PVA containing brain organoids were related to axon and synapse maturation (*GREM1, FABP5, GAREM2, MAP6D1, ND1, ND2, RAB26, FLOT1*), with many of these genes relating specifically to glutaminergic neuron maturation (*GREM1, FABP5, ND1, ND2, FLOT1*). Light-trained samples also exhibited downregulation of neuronal migration genes (*RSPH3, DSEL, PLPR3*), which are usually heightened in immature neurons during development,^29,30^ and upregulation of genes related to eye and retina development (*ALDH3A1, FJX1, RDH13, RARRES2, PRPH2, PFDN5, RGR*) (Fig. 5d-e and Table S5). Many DEGs in the light-trained, PVA-only control organoids (-rGO +Light) also relate to neuronal and retinal maturation, but expression is often higher with a combination of light-training effects and rGO-PVA nanofiber effects (Fig. S7).

Gene ontology (GO) enrichment analysis of DEGs between light-trained and unstimulated brain organoids with rGO-PVA nanofibers indicated strong enrichment in retinal, immune response, and oxidative phosphorylation gene transcription in response to light stimulation (Fig. 5f-g and Fig. S8a-c) and a slight reduction in forebrain development gene transcription (Fig. 5g), perhaps as a direct result of the increase in retinal gene expression. Interestingly, GO analysis also indicated strong enrichment of ATP synthesis and oxidative phosphorylation, which is required for neuronal maturation^31^, in unstimulated brain organoids with rGO-PVA nanofibers compared to PVA-only control (Fig. S8c-d), suggesting some degree of neuronal maturation occurs due to the presence of rGO, even in the absence of light stimulation. Gene Set Enrichment Analysis (GSEA) further confirmed these trends, including increases in ATP synthesis and decreases in GABAergic pathways in light-trained organoids with rGO-PVA nanofibers (Fig. S8e).

### 3.5 Retinal cellular structures, more excitatory neurons, and fewer forebrain neurons are found in light-trained 3D brain organoids with rGO-PVA nanofibers

Next, we imaged proteins indicative of neural and retinal maturation in brain organoids with nanofibers to confirm transcriptomic results. We found that light-trained brain organoids with rGO-PVA nanofibers had more cells stained positive for the retinal markers (S-OPSIN+, CRX+, RCVRN+ and RHO+ cells) (Fig. 6a and Fig. S9a). Co-localized clusters of CRX+ and RCVRN+ cells, which mark early photoreceptor cell formation,^32,33^ are visible in both light-trained and unstimulated brain organoids (Fig. S9a), although they are not localized to the organoids’ outer peripheries as in conventionally differentiated retinal organoids^34^. In contrast, light-trained brain organoids with rGO-PVA nanofibers uniquely exhibit clusters of S-OPSIN-rich cone cells along the organoid surface, presumably where more light penetrated the organoid. There was no presence of a retinal pigment epithelium in any condition, i.e., there was a lack of ZO-1+ cells. S-OPSIN+ cells were surrounded by MAP2+ neurons, which may include retinal ganglion cells that are responsible for receiving light signals from photoreceptor cells as electrical signals^34,35^ (Fig. 6a). The greater numbers of MAP2+ neurons in the light-trained brain organoids with rGO-PVA nanofibers may contribute to heightened light-responsiveness in MEA measurements. Since RNA analysis indicated decreased forebrain development (Fig. 5g), we also characterized proteins related to forebrain maturity. Light-trained brain organoids with rGO-PVA nanofibers had fewer DCX+ migratory neurons as well as lower concentrations of FOXG1 proteins and fewer FOXG1+ forebrain cells (Fig. 6b). DCX+ cells were restricted to the outermost surface of brain organoids with rGO-PVA nanofibers, though they appear more pervasive in both unstimulated brain organoids with rGO-PVA nanofibers and light-trained brain organoids with PVA-only fibers (Fig. 6b and Fig. S9b), suggesting that light-activation of rGO impairs forebrain development as opposed to the presence of light or rGO alone. Together, these findings indicate light-trained organoids with rGO-PVA nanofibers exhibit impaired forebrain differentiation and cortical neuron migration as they shift towards retinal cell differentiation.

**Fig. 6.**
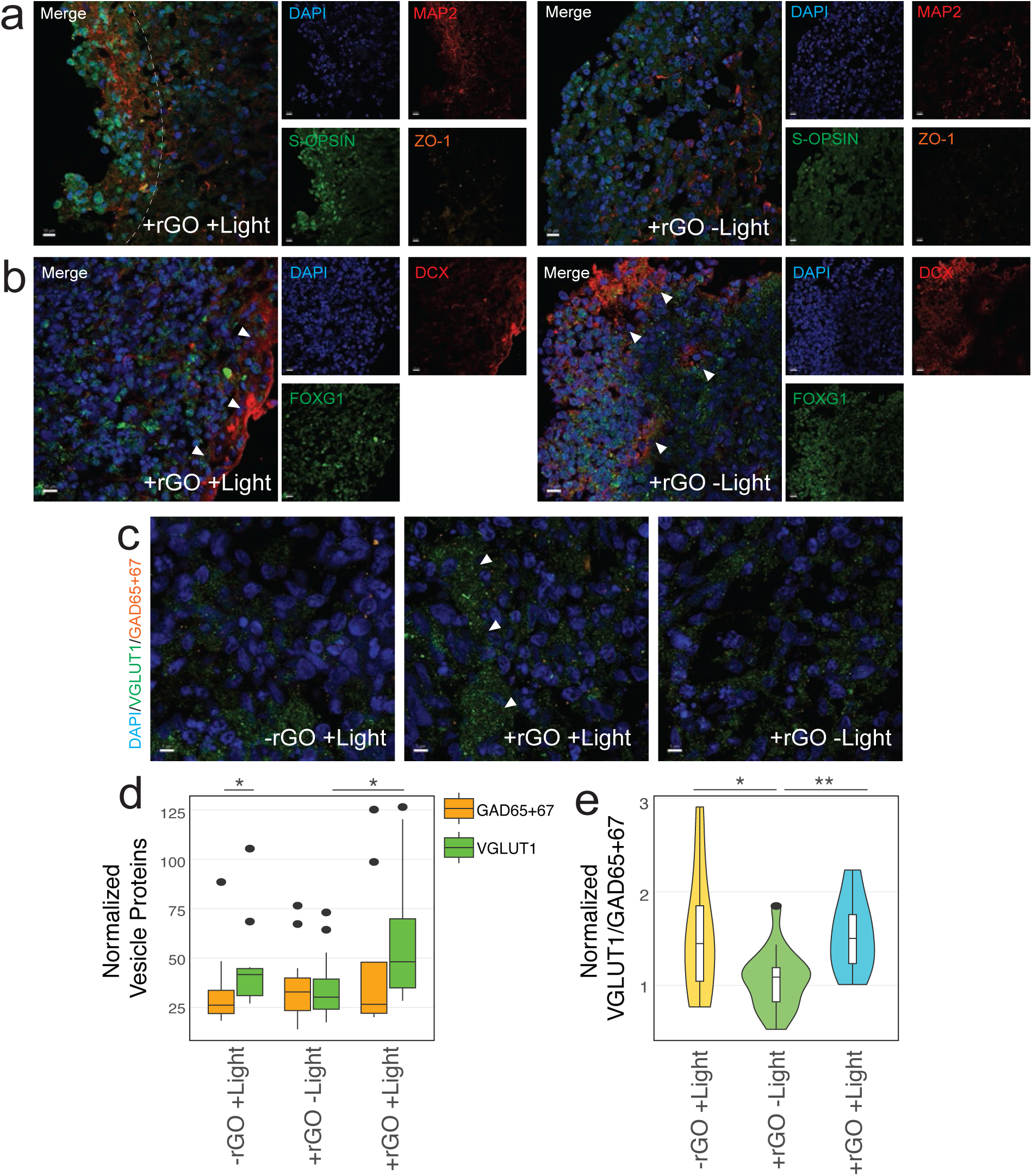
Fluorescent images of light-trained brain organoids with rGO-PVA nanofibers show enhanced retinal cell differentiation, fewer migrating neurons, and more glutaminergic vesicle proteins. Images of brain organoids with nanofibers after 8 weeks of light-training on MEAs stained for (**a)** MAP2 (red), S-OPSIN (green), ZO-1 (orange) and (**b)** DCX (red) and FOXG1 (green). The dashed line in panel a indicates regions higher in MAP2+ (left of line) and S-OPSIN+ cell layers (right of line). Arrows in b point to the boarder of DCX+ cells. Scale bar = 10 µm. **(c)** Images of brain organoids with nanofibers after 8 weeks of light-training on MEAs stained for VGLUT1 (green) and GAD65/67 (orange). Arrows point to clusters of VGLUT1+ puncta. Scale bar = 5 µm. **(d)** Quantification of VGLUT1+ and GAD65+67+ puncta normalized to DAPI+ cells in brain organoids in c. (N = 2-3 organoids per condition and batch; n = 11, 16, and 8 ROIs per cell). Median values of -rGO +Light, +rGO -Light, and +rGO +Light were 41.6, 30.2, and 48.1, for VGLUT1+ and 26.2, 32.8, and 26.6, for GAD65+67+, respectively. Kruskal-Wallis, p=0.063. **(e)** Quantification of VGLUT1+ and GAD65+67+ puncta normalized to DAPI+ cells in brain organoids in c as a ratio. (N = 2-3 organoids per condition; n = 11, 16, and 8 ROIs). Median values of -rGO +Light, +rGO -Light, and +rGO +Light were 1.45, 1.09, and 1.50, respectively. Kruskal-Wallis, p=0.016. Statistical analyses were made using Kruskal-Wallis and Wilcoxon Tests and significance is defined as * p<0.5, ** p<0.01, *** p<0.001, and **** p<0.0001.

Lastly, we examined synaptic vesicle proteins to evaluate if there is greater excitatory neurotransmission in light-trained organoids with rGO-PVA nanofibers. In all organoids with rGO-PVA nanofibers, localization of vesicular glutamate transporter family member 1 (VGLUT1), a glutaminergic synaptic vesicle protein, and glutamic acid decarboxylases 65 and 67 (GAD65+67), proteins that synthesize and package GABA into vesicles,^36^ appear co-localized in highly concentrated clusters (Fig. S9c). Outside of these regions, GAD65+67+ puncta were about evenly distributed in all conditions. However, VGLUT1 distribution varied, with dense clusters of VGLUT1+ puncta visible in light-trained organoids with rGO-PVA nanofibers (Fig. 6c and Fig. S9c). Both synaptic vesicle protein types were equally present in unstimulated brain organoids, but in both light-stimulated organoids with rGO-PVA fibers and PVA-only fibers, GAD65+67+ puncta were slightly decreased while VGLUT1+ puncta significantly increased (Fig. 6c-e). These results indicate that repetitive light-training in brain organoids with nanofibers increases excitable neuronal differentiation and the expression of proteins required for excitable functions.

## 4. Conclusion and Discussion

Utilizing hPSC-derived EECs for disease modeling and developmental studies remains particularly challenging given their suboptimal electrical maturity compared to *in vivo* cells.^1–6^ Here, we implement PVA-based electrospun nanofibers containing rGO to impact electrical action potentials in hPSC-derived cardiomyocytes and neurons. Through light-activation of rGO-PVA nanofibers, we demonstrated their ability to improve the electrical activity in hPSC-EECs in the presence of rGO and in response to acute light stimulation. Chronic, daily light training over 8 weeks enhanced the electrical response of brain organoids to light by maturing excitatory neurons and driving retinal cell differentiation. Despite these benefits to electrical maturity, we would like to provide three caveats and context for these data.

First, rGO-PVA can be electrospun into nanofibers that have stable mechanical and electrical properties similar to other conductive fibers.^7,15,37–39^ However, nanofiber scaffolds with higher rGO content (0.01-0.1% rGO) were challenging to fabricate due to the enhanced hydrophobic separation of their nanocomposites,^14,15^ leading to increasingly variable fiber density, fiber diameter, and electrical conductivity (Fig. 1). The latter is of particular importance, since conductivity is reflective of rGO content and varies with greater electrical and light-responsiveness in EECs. Thus, exploring strategies to reliably and uniformly increase rGO content in nanofibers is imperative. Techniques like hot-pressing may homogenize rGO-PVA nanocomposites for longer durations prior to electrospinning,^40^ enabling more even rGO distribution throughout fibers. Nanocomposites of graphene and polymers with moderate hydrophobicity like polycaprolactone (PCL) and polyethylene terephthalate (PET) have successfully yielded high graphene content electrospun fibers,^15,37–39^ but typically require greater chemical functionalization to promote cellular attachment than hydrophilic polymers like PVA.^41–44^ Alternatively, some studies have deposited graphene onto electrospun polymer nanofibers,^16,45^ which may facilitate greater graphene-cell interactions since here, flakes are not embedded in the fibers.

Second, cell attachment to nanofibers was significantly enhanced by coating the fibers with Matrigel. Although scaffold characterization revealed that Matrigel-coating reduced the conductivity of rGO-PVA scaffolds (Fig. S3c), this was not reflected in reduced cellular electrical performances during functional assays, which consistently showed that electrical activity varies with rGO concentration (Fig. 2h, 3g, 4f-g and Fig. S1d, S2e, S4b, S5). Still, cell attachment was moderately variable, leading to irregular cell densities on 2D scaffolds that may have limited endogenous electrical activity and cellular responsiveness to the material. Notably, rGO-PVA nanofibers saw enhanced cell attachment compared to PVA-only nanofibers, likely due to the beneficial effects of surface charge on cellular adhesion.^46^ Electrospinning aligned fibers may promote greater cardiomyocyte adhesion, proliferation, and migration versus randomly aligned fibers as shown in other fiber system.^47–51^ Neurons, on the other hand, readily attached to our fibers and received signals from them, mirroring the dynamic relationship of neurons and the extracellular matrix during *in vivo* development.^52,53^ On MEAs, fiber rearrangement introduces the possibility that they could obstruct cell-electrode connections, reducing the number of action potentials detected by MEA. Given that fiber attachment is a critical aspect of whether cell function improves, the possibility that some variance arose from adhesion differences is worth noting.

We believe that the degree of retinal cell development in light-trained brain organoids with rGO-PVA nanofibers is a unique discovery resulting from the combination of chronic, repetitive light exposure and electrical stimulation from the rGO nanofibers. We suspect transcript signals of retinal markers were dilute–and retinal structures were localized to the outside of the organoids (Fig. 6a and Fig. S9a)–because nanofiber and light interactions may have been limited to the organoid’s surface. We hypothesize that the development of retinal cells in light trained organoids with rGO-PVA nanofibers contributed to heightened electrical responses over time, because light and rGO can photosensitize cells so that they can detect and transmit light signals to electrical signals.^54,55^ However, the origins of these photosensitive cells remains unclear. Since we employed older brain organoids for these experiments, these data suggest plasticity in neuronal fates and circuitry in response to environmental factors,^56–59^ perhaps enabling the transition from mature neurons into photoreceptor cells. Alternatively, retinal cell populations are known to exist in older brain organoids,^24^ so light stimulation may instead expand already existing retinal progenitor populations and promote their localization to organoid surface layers.

Finally, the cause of heightened retinal cell differentiation is also yet to be elucidated, but we propose several potential contributing factors. First, external sensory stimulation is known to play key roles in the development of sensory circuits and tissues *in vivo*.^54,60^ Mammals are born with immature photoreceptors that undergo rapid maturation on genetic and cellular levels in response to light shortly after birth.^54,61^ In this study, light stimulation, enhanced by matched electrical pulses from activated rGO, may mimic these important developmental events promoting retinal maturation. We additionally suggest that the presence of environmental stresses in our cell culture system–incidentally derived from repetitive stimulation–may also contribute to retinal cell differentiation. One potential source of oxidative stress is via the oxidation of vitamin A in the media by free radicals produced by rGO activation^62–64^ or UV light.^65–67^ The byproduct, retinal, is reduced to the pro-retinal differentiation metabolite retinoic acid,^68,69^ by enzymes like retinol dehydrogenase 13 (*RDH13*) and aldehyde dehydrogenase 3A1 (*ALDH3A1)*, two of the most significantly elevated DEGs in light-trained organoids with rGO-PVA nanofibers (Fig. 5d, e, and Fig. S7). Another potential source of environmental stress may be heat discharged as a byproduct of light stimulation. Regardless of this surprising finding, clearly further mechanistic investigation of retinal cell differentiation via chronic light-activation of rGO-PVA nanofibers in needed.

With this context, light-activation of rGO-PVA nanofibers appears to provide a robust, novel method to stimulate electrical activity in hPSC-derived EECs both on-demand and over time. Combining rGO nanofibers with hPSC-derived cells brings us closer to having more completely electrically mature and functionally accurate *in vitro* tissues that may better mimic human development as well as disease pathways. This study also suggests that the importance of sensory input in *in vitro* cell cultures may be less well considered; minimally controlled factors in cell culture like light and sound may play a larger role in stem cell maintenance and differentiation than previously thought. On the other hand, sensory inputs may be applied to accelerate certain cell type specifications, as exemplified here by the development of retinal cells in 8 weeks, which is notably faster than using traditional biomolecule-based retinal organoid differentiations.^34,70,71^ These hybrid organoids may accomplish a level of connectivity that is strived for in cortex/retinal assembloids, without the need to perform multiple, complex differentiations and challenging techniques to combine organoids.^35,72^ Given these advances with light-reactive rGO-PVA fibers, we believe that this study presents a new biomaterial to further mature hPSC-derived EECs.

## Acknowledgements

The authors thank Dr. Kristen Jepsen and the Institute for Genomic Medicine (UCSD) for preparing and sequencing RNA libraries and the Nanotechnology Infrastructure at Nano3 (UCSD), part of the National Technology Coordinated Infrastructure funded by the National Institutes of Health (ECCS-2025752), for access to and maintenance of the Dektak XT Stylus Profilometer, Zeiss Sigma 500, Emitech Sputter Coater, and Disco Automatic Dicing Saw 3220. We would also like to thank Dr. Alex Savchenko for providing the rGO flakes and PhotonMaker device. In addition, we would like to thank the Stem Cell Genomics Core at the Sanford Stem Cell Institute for providing access to and maintenance of the Dragonfly 600 microscope. This publication includes data generated at the UC San Diego IGM Genomics Center utilizing an Illumina X Plus that was purchased with funding from a National Institutes of Health SIG grant (#S10 OD026929). The authors acknowledge funding support from the National Institutes of Health (R01NS116802 to A.E. and 1R01MH128365, R01NS123642, 1R01ES033636, MH123828, MH127077, and NS105969 to A.R.M.), the NSF Graduate Research Fellowship Program (to E.L.), and CIRM Training Program (EDUC4-12804 to E.L.) for additional fellowship support.

## Footnotes

### CRediT Authorship Contribution Statement

**Erin LaMontagne:** Conceptualization, Investigation, Writing – original draft, Visualization. **Gisselle Gonzalez:** Resources, Methodology. **Ritwik Vatsyayan:** Methodology, Investigation. **Blanca Martin-Burgos:** Methodology, Investigation. **Francesca Puppo:** Methodology. **Diogo Biagi:** Methodology. **Fabio Papes:** Methodology. **Shadi A. Dayeh:** Writing – review & editing. **Alysson R. Muotri:** Writing – review & editing, Supervision, Conceptualization, Funding acquisition. **Adam J. Engler:** Writing – original draft, Supervision, Conceptualization, Funding acquisition.

### Data Availability

RNA sequencing results are accessible through NCBI via Gene Omnibus Express number GSE26898. All other data is available upon request. Software to analyze calcium imaging as well as bulk RNA-sequencing are available via Github. (https://github.com/eelamonta/Graphene-Project). Other data will be made available on request.

---

**Supplemental Fig. 1.**
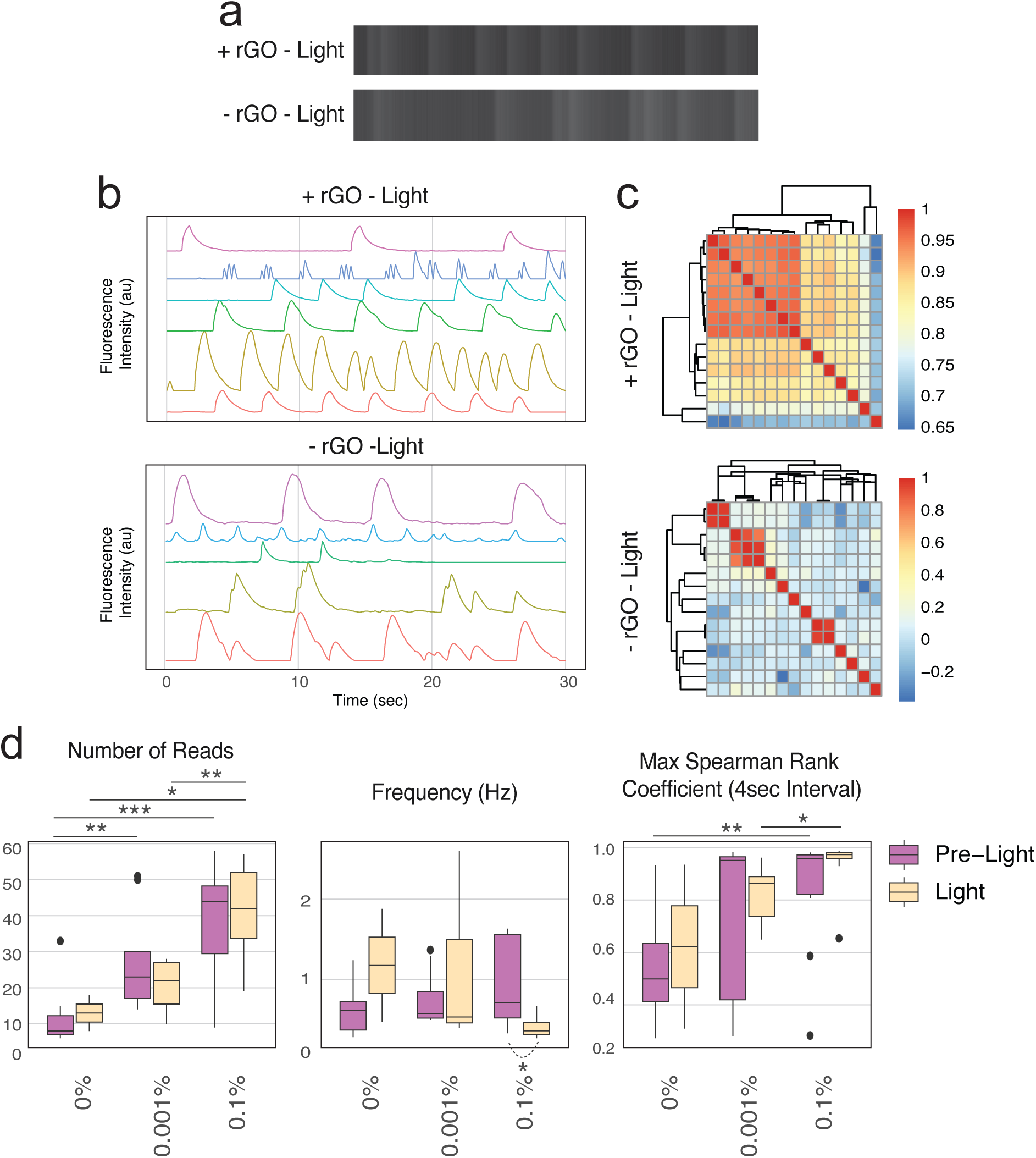
Cardiomyocytes on rGO-PVA scaffolds without light stimulation are more synchronous. **(a)** Representative kymographs, **(b)** light intensity profiles, and **(c)** heatmaps of Spearman Rank Coefficients of cardiomyocytes on scaffolds with and without rGO, without light stimulation. **(d)** Calcium handling metrics of cardiomyocytes on scaffolds containing 0%, 0.001%, and 0.1% rGO with and without acute light stimulation. Number of reads, frequency (Hz) and max Spearman Rank Coefficient (4 sec interval) have Kruskal-Wallis p-values of 0.00028, 0.11, and 0.0042, respectively. Median values are reported in Table S1. (N = 4-5 cardiomyocyte differentiation batches; n = 10, 9, 12 for 0%, 0.001%, and 0.1% rGO without light, respectively; n = 2, 7, 8 for 0% 0.001%, and 0.1% rGO with light, respectively). Statistical analyses were made using Kruskal-Wallis and Wilcoxon Tests and significance is defined as * p<0.5, ** p<0.01, *** p<0.001, and **** p<0.0001.

**Supplemental Fig. 2.**
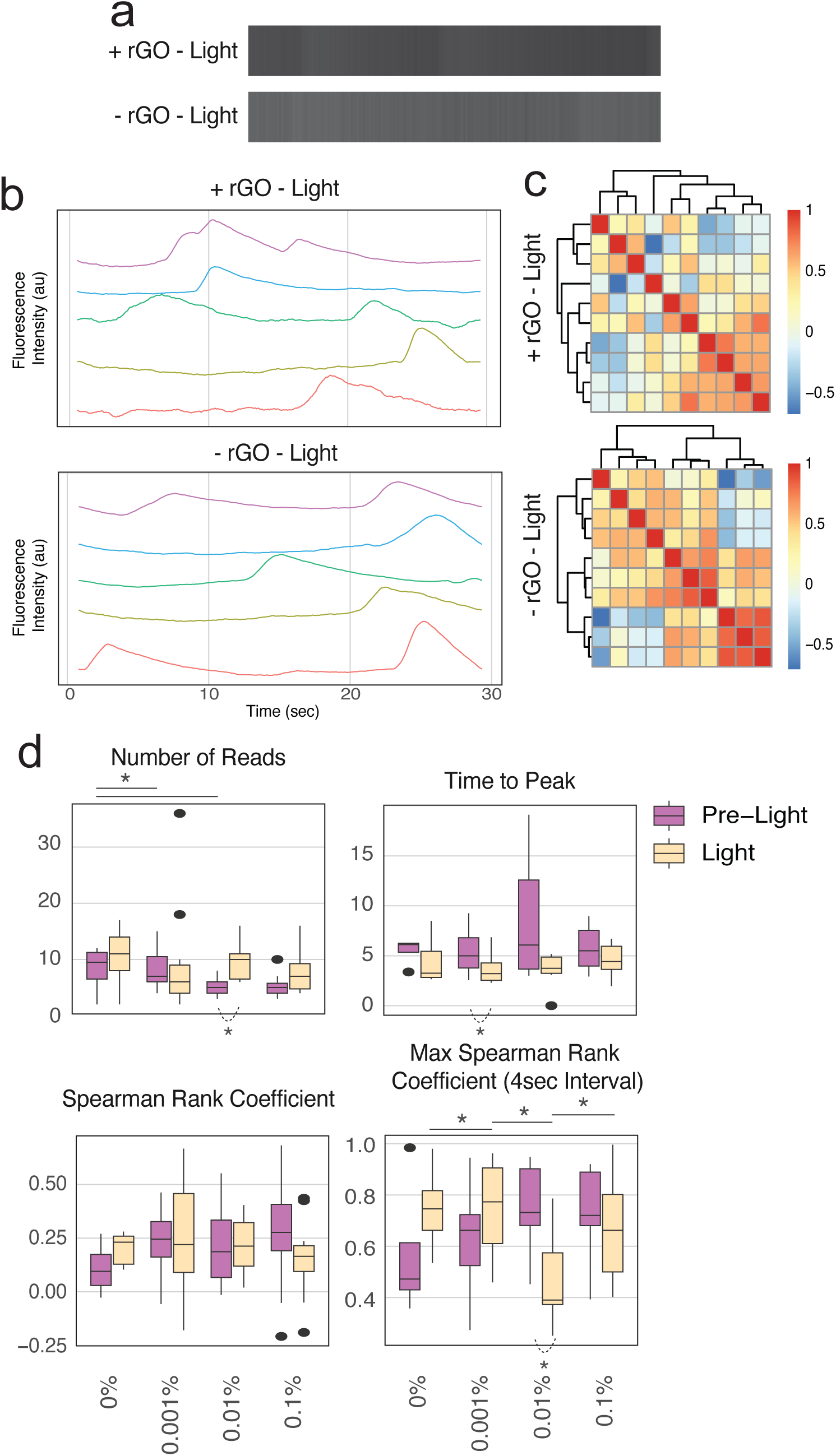
Neurons on rGO-PVA scaffolds without light stimulation are functionally similar to neurons on PVA-only scaffolds. **(a)** Representative kymographs, **(b)** high temporal resolution calcium trace with progressive smoothing by computational processing from (i) a raw trace, (ii) the raw trace smoothed via moving average, (iii) the smoothed trace with labeled local maxima (calcium event peaks; red dots) and local minima (blue dots) used to determine the signal baseline (blue line), and (iv) the smoothed, baseline-corrected calcium trace. **(c)** representative light intensity profiles, and **(d)** heatmaps of Spearman Rank Coefficients of neurons on scaffolds with and without rGO, without light stimulation. **(e)** Calcium handling metrics of neurons on scaffolds containing 0%, 0.001%, 0.01%, and 0.1% rGO with and without acute light stimulation. Number of reads, time to peak, Spearman Rank Coefficient and max Spearman Rank Coefficient (4 sec interval) have Kruskal-Wallis p-values of 0.1, 0.085, 0.77, and 0.077, respectively. Median values are reported in Table S2. (N = 5 brain organnoid differentiation batches; n = 4, 9, 12, 10 for 0% 0.001%, 0.01%, and 0.1% rGO without light, respectively; n = 6, 11, 7, 12 for 0%,0.001%, 0.01%, and 0.1% rGO with light, respectively). Statistical analyses were made using Kruskal-Wallis and Wilcoxon Tests and significance is defined as * p<0.5, ** p<0.01, *** p<0.001, and **** p<0.0001.

**Supplemental Fig. 3.**
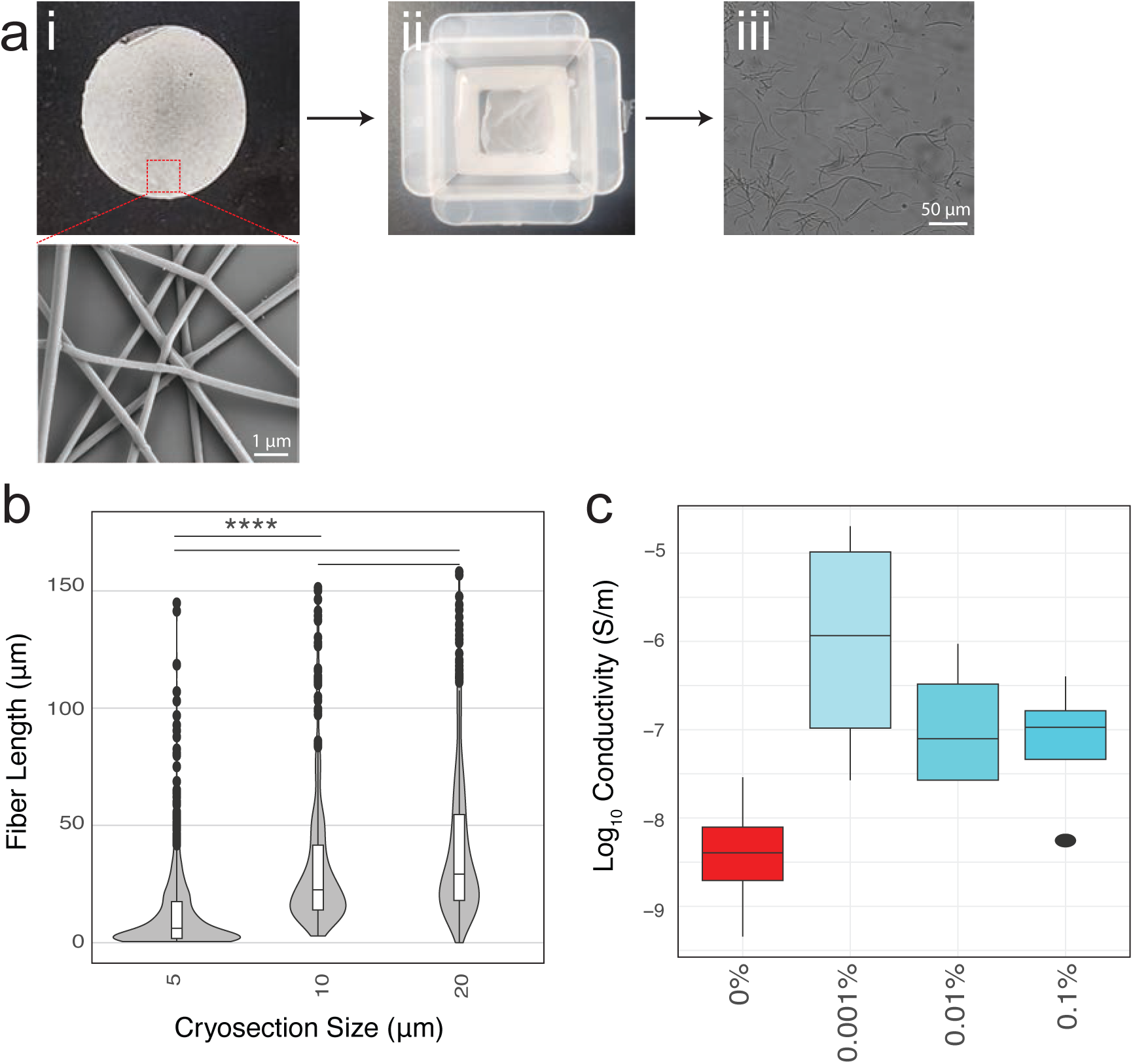
Nanofiber fabrication and Matrigel-coated scaffold conductivity. **(a)** Processing of electrospun scaffolds into a nanofiber suspension by electrospinning scaffolds on glass coverslips visualized by (i) eye and SEM (Magnification = 15,000 X, Scale bar = 1 µm). (ii) Crosslinked scaffolds are frozen, cryosectioned, and filtered. (iii) Nanofibers are lyophilized, resuspended, and visualized by light microscopy (Scale bar = 50 µm). **(b)** Nanofiber length measurements from light microscopy images (n = 500 measurements per cryosection size). Median nanofiber lengths of scaffolds cryosection at 5, 10, and 20 µm were 6.13, 23.0, and 29.8 µm, respectively. Kruskal-Wallis, p<2.2e-16. **(c)** Conductivity of Matrigel-coated scaffolds (N = 4 scaffolds per condition, n = average of 6-8 measurements per scaffold). Median conductivities of 10% PVA scaffolds containing 0%, 0.001%, 0.01%, and 0.1% rGO were 4.15e-9, 4.21e-6, 1.30e-7, and 1.07e-7 S/m, respectively. Kruskal-Wallis, p=0.079. Statistical analyses were made using Kruskal-Wallis and Wilcoxon Tests and significance is defined as * p<0.5, ** p<0.01, *** p<0.001, and **** p<0.0001.

**Supplemental Fig. 4.**
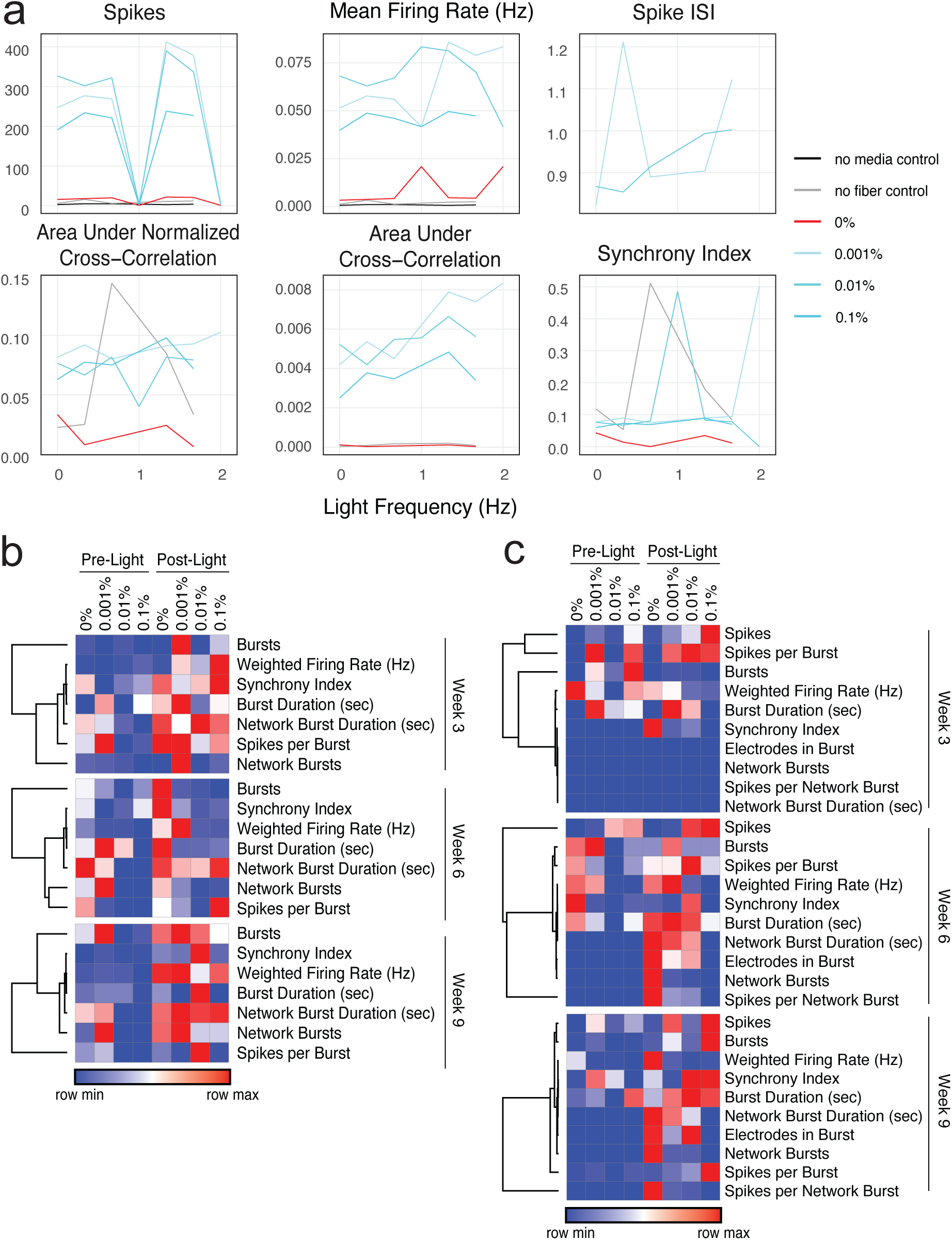
Data showing fiber-only controls and supplemental brain organoid with rGO-PVA nanofibers MEA metrics in response to acute light stimulation. **(a)** MEA metrics produced by an empty well, a well with cell culture media, and wells with culture media and nanofibers during light stimulation at various frequencies (n = 1 MEA well per condition). Metrics reported are the only metrics altered by nanofibers. **(b)** Heatmaps of median values of MEA metrics produced by brain organoids at 3-, 6-, and 9-weeks post-fiber addition, comparing pre- and post-light recordings. Complementary data to Fig. 4f. (N = 3 brain organoid differentiation batches; n = 36-45 MEA wells per condition). MEA metric values are reported in Table S3.

**Supplemental Fig. 5.**
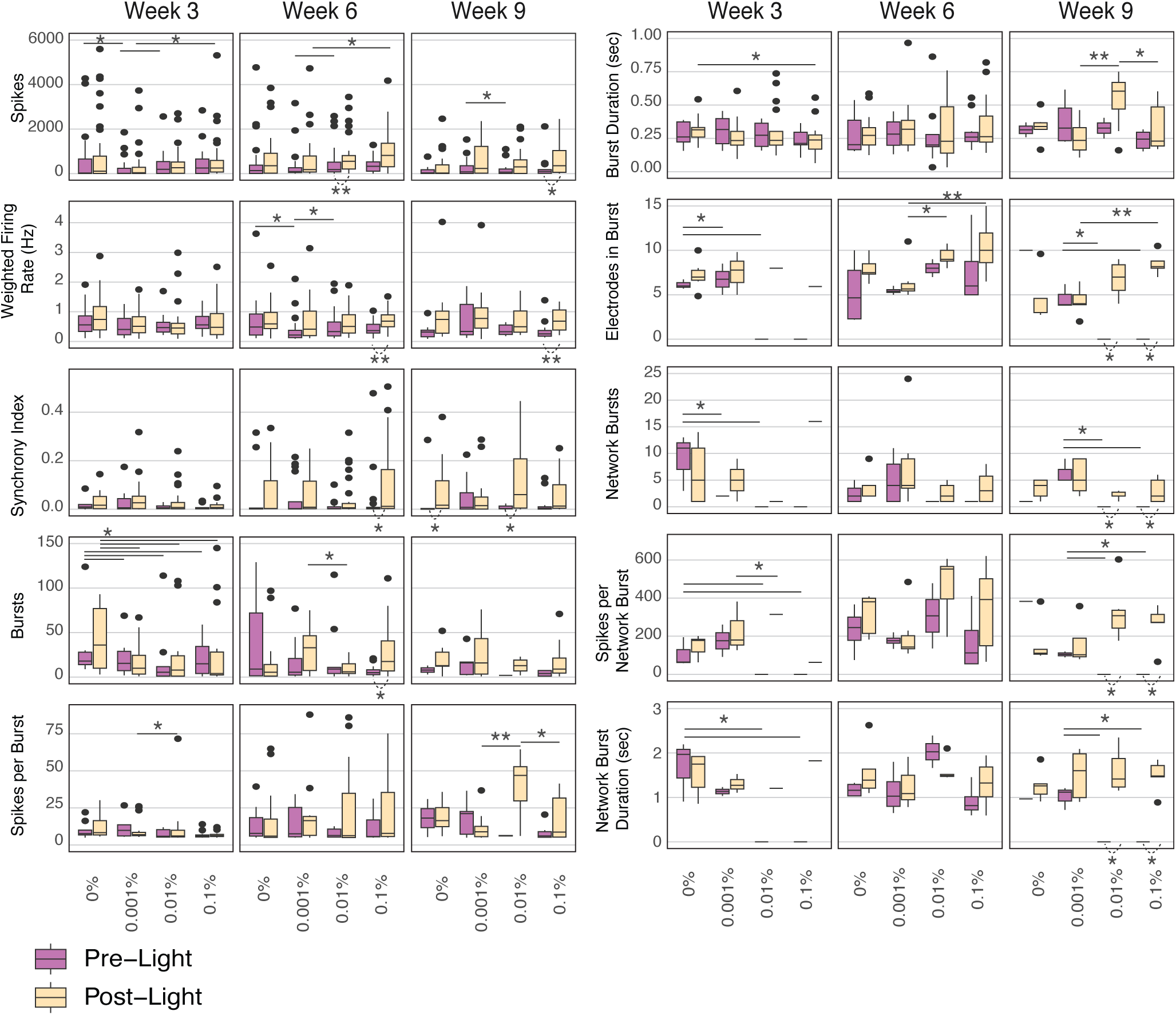
Brain organoids with rGO-PVA nanofibers MEA metrics in response to acute light stimulation over multiple weeks. MEA metrics of brain organoids with nanofibers before and after acute light stimulation during 8 weeks of daily, chronic light training at 3-, 6-, and 9-weeks post-fiber addition (N = 3 brain organoid differentiation batches; n = 36-45 MEA wells per condition). MEA metric values are reported in Table S3 and Kruskal-Wallis p-values are reported in Table S4. Statistical analyses were made using Kruskal-Wallis and Wilcoxon Tests and significance is defined as * p<0.5, ** p<0.01, *** p<0.001, and **** p<0.0001.

**Supplemental Fig. 6.**
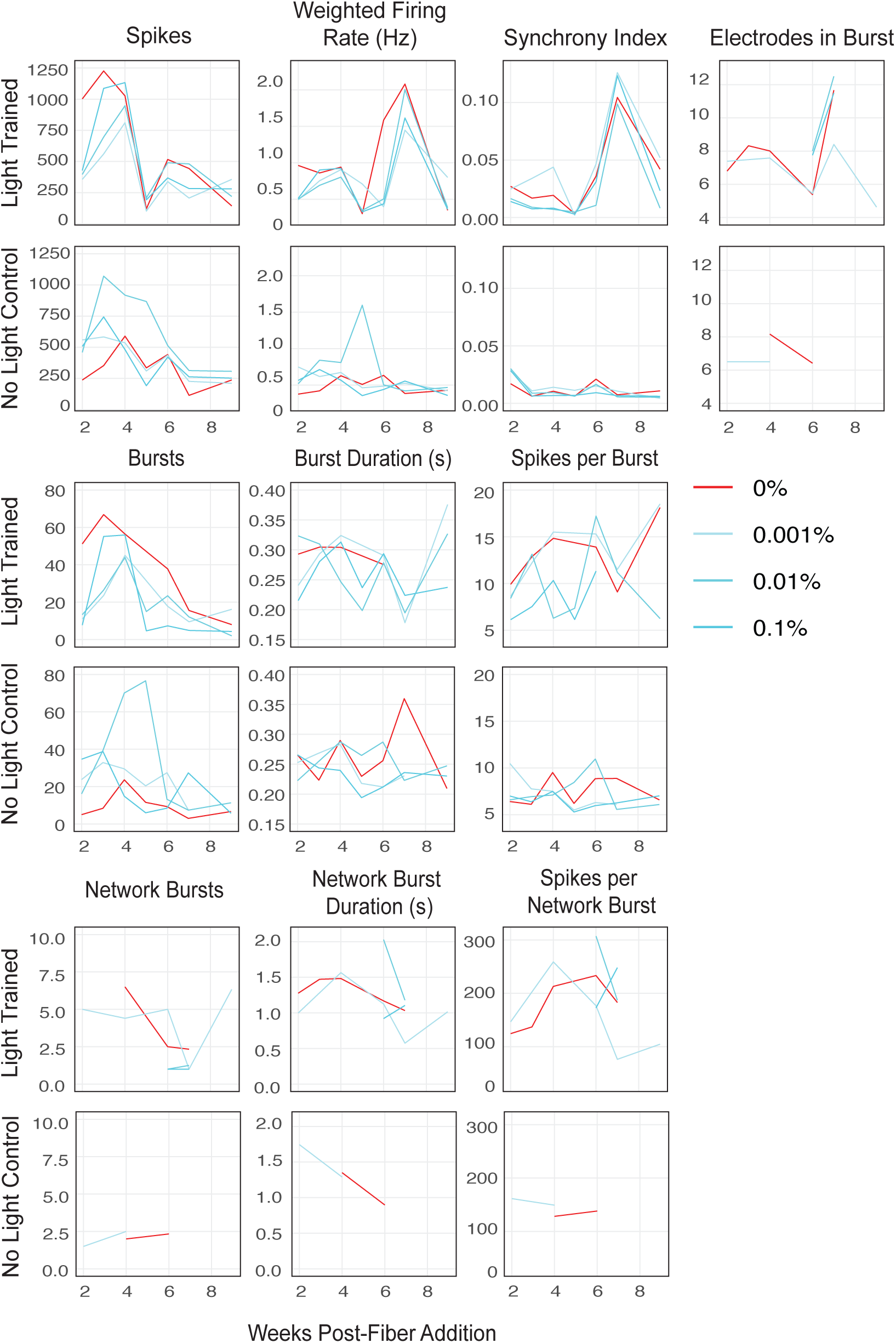
Function of light-trained brain organoid compared to unstimulated brain organoids. Mean MEA metrics of brain organoids with nanofibers pre-light stimulation during 8 weeks of daily, chronic light training compared to brain organoids with nanofibers never exposed to light stimulation during the 8 weeks. (N = 3 brain organoid differentiation batches; n = 36-45 MEA wells per condition). MEA metric values are reported in Table S3.

**Supplemental Fig. 7:**
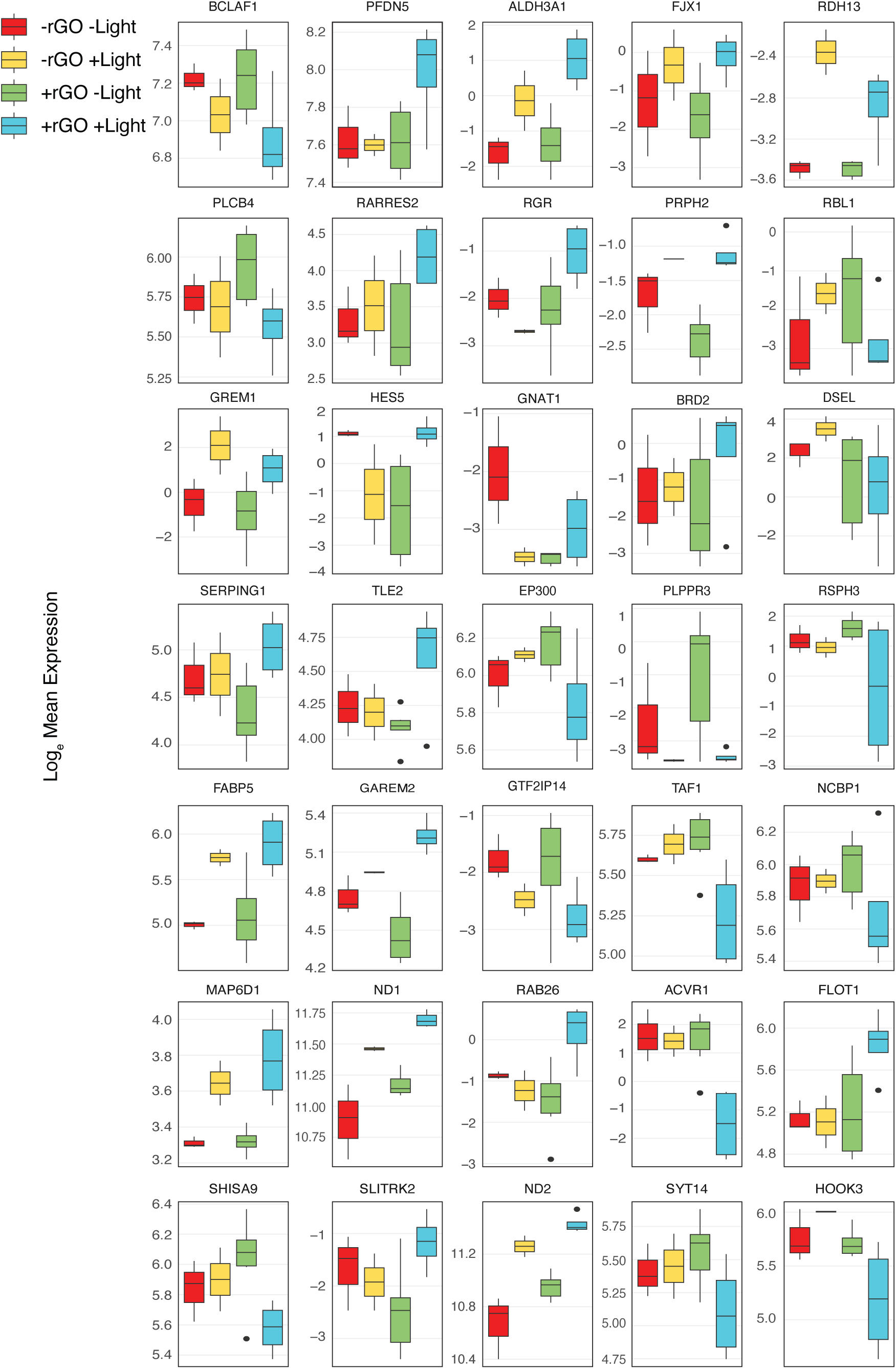
Expression of significant DEGs in light-trained brain organoid compared to unstimulated brain organoids. Boxplots of log_e_ mean expression of significant DEGs between light-trained and unstimulated brain organoids with rGO-PVA nanofibers (significant DEGs are described in Fig. 5d). Normalized transcript counts are reported in Table S5.

**Supplemental Fig. 8:**
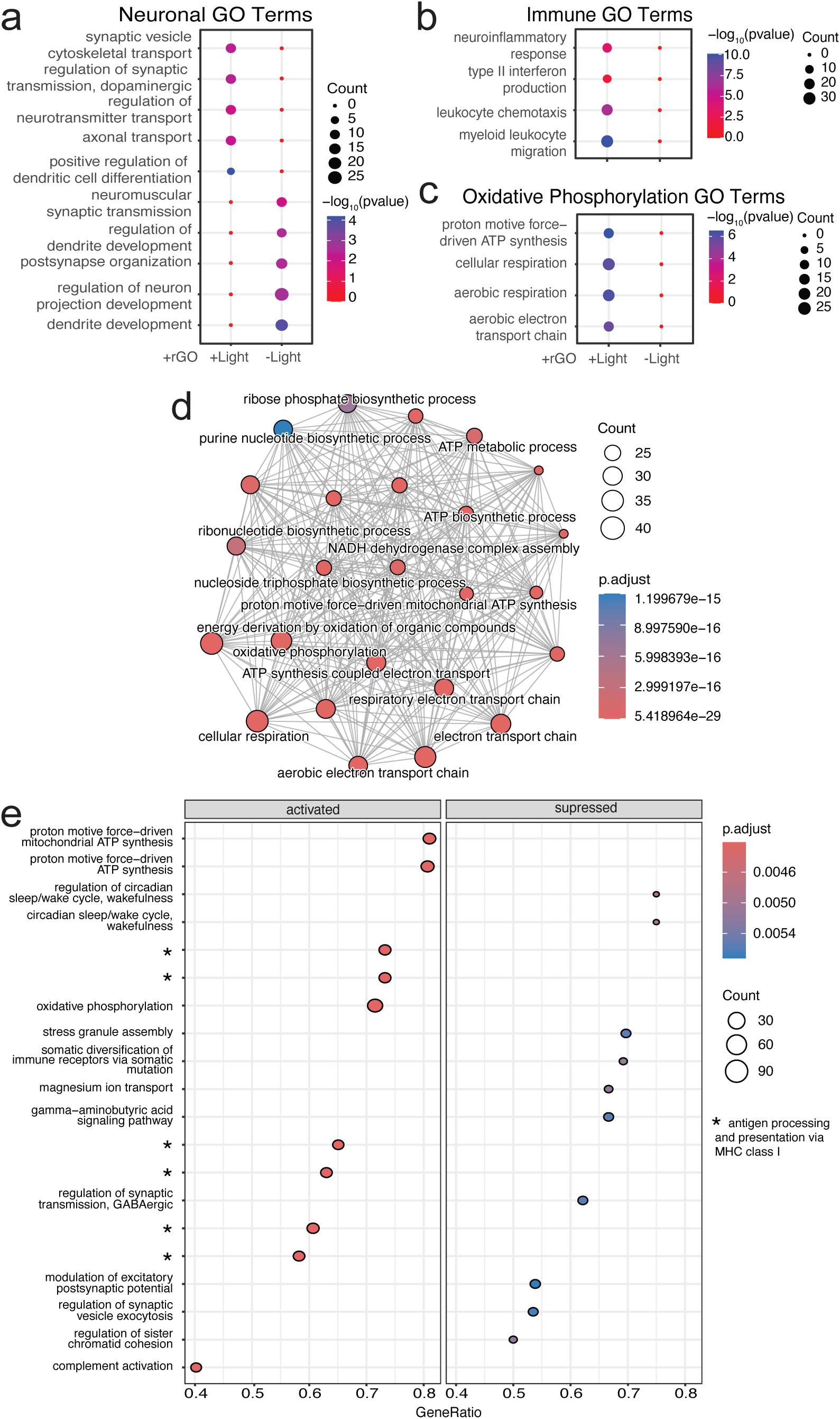
Supplemental GO term and GSEA analysis of light-trained brain organoid compared to unstimulated brain organoids. Enriched GO terms from DEGs between light-trained and unstimulated brain organoids with rGO-PVA nanofibers related to **(a)** neuronal maturation and synapse formation, **(b)** immune functions, and **(c)** oxidative phosphorylation. **(d)** Emap of top 25 DEGs between unstimulated organoids with rGO-PVA nanofibers versus light-trained and unstimulated organoids with PVA-only fibers. **(e)** GSEA pathways activated and suppressed in light-trained organoids with rGO-PVA nanofibers verses unstimulated organoids with rGO-PVA nanofibers. Normalized transcript counts are reported in Table S5.

**Supplemental Fig. 9:**
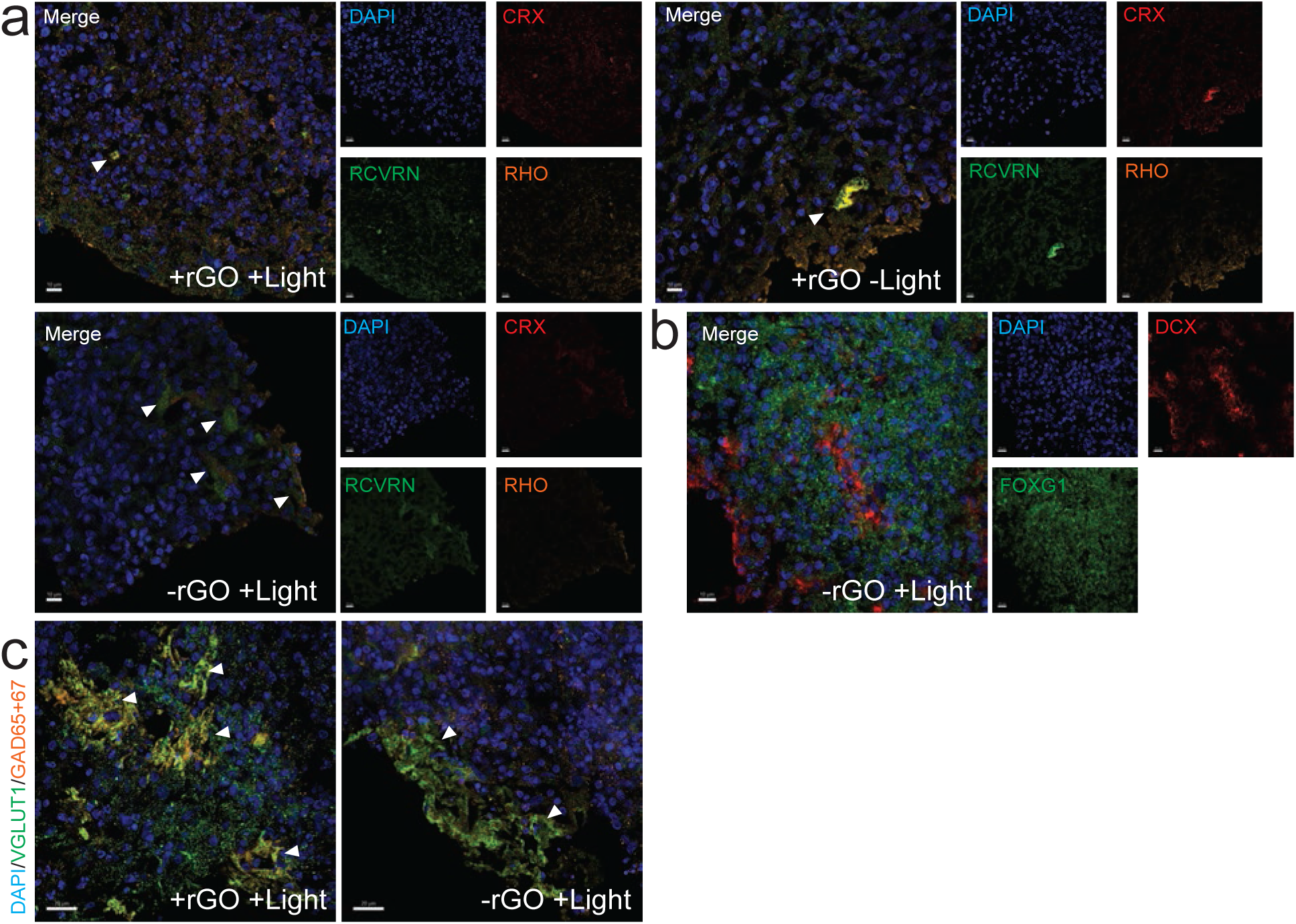
Supplemental fluorescent images of light-trained brain organoids with rGO-PVA nanofibers. Images of brain organoids with nanofibers after 8 weeks of light-training on MEAs stained for **(a)** CRX (red), RCVRN (green), RHO (orange) and **(b)** DCX (red) and FOXG1 (green). Arrows in a point to clusters of RCVRN+ and CRX+ cells. Scale bar = 10 µm. **(c)** Images of brain organoids with nanofibers after 8 weeks of light-training on MEAs stained for VGLUT1 (green) and GAD65+67 (orange). Arrows point to clusters of vGLUT1/GAD65+67+ puncta. Scale bar = 20 µm.

## Supplemental Tables

**Supplemental Table 1. Calcium handling metrics of cardiomyocytes on scaffolds.** Table of statistical summaries for calcium handling metrics of cardiomyocytes in Fig. 2h and Fig. S1d. The number of reads equates to the number of cells analyzed per biological replicate.

**Supplemental Table 2. Calcium handling metrics of neurons on scaffolds.** Table of statistical summaries for calcium handling metrics of neurons in Fig. 3g and Fig. S2d. The number of reads equates to the number of cells analyzed per biological replicate.

**Supplemental Table 3. MEA metrics of brain organoids with nanofibers in response to acute light stimulation during daily, chronic light training.** Table showing statistical summaries for the MEA metrics of brain organoids at 3-, 6-, and 9-weeks post-fiber addition visualized in Fig. 4f-g, and Fig. S4b, and S5.

**Supplemental Table 4. Significance of MEA metrics by Kruskal-Wallis.** Table showing Kruskal-Wallis p-values for MEA metrics of brain organoids at 3-, 6-, and 9-weeks post-fiber addition visualized in Fig. 4f and Fig. S5.

**Supplemental Table 5. RNA sequencing expression matrix.** Table showing the normalized log_2_ expression value for each gene (rows are gene Ensembl IDs) and technical replicate (columns are “batch number (arbitrary)_rGO concentration_light or no light treatment”). This data was used to derive and analyze DEGs and expression visuals in Fig. 5 and Fig. S7, S8.

## References

1. Risner-Janiczek, J. R., Ungless, M. A. & Li, M. Electrophysiological Properties of Embryonic Stem Cell-Derived Neurons. PLoS ONE 6, e24169 (2011).

2. Chan, Y.-C. et al. Electrical Stimulation Promotes Maturation of Cardiomyocytes Derived from Human Embryonic Stem Cells. J. of Cardiovasc. Trans. Res. 6, 989–999 (2013).

3. Puppo, F. et al. All-Optical Electrophysiology in hiPSC-Derived Neurons With Synthetic Voltage Sensors. Front. Cell. Neurosci. 15, 671549 (2021).

4. Sakamoto, K. et al. Emergent synchronous beating behavior in spontaneous beating cardiomyocyte clusters. Sci Rep 11, 11869 (2021).

5. Halliwell, R. F., Salmanzadeh, H., Coyne, L. & Cao, W. S. An Electrophysiological and Pharmacological Study of the Properties of Human iPSC-Derived Neurons for Drug Discovery. Cells 10, 1953 (2021).

6. Trujillo, C. A. et al. Complex Oscillatory Waves Emerging from Cortical Organoids Model Early Human Brain Network Development. Cell Stem Cell 25, 558–569.e7 (2019).

7. Gonzalez, G. et al. Conductive electrospun polymer improves stem cell-derived cardiomyocyte function and maturation. Biomaterials 302, 122363 (2023).

8. Savchenko, A. et al. Graphene biointerfaces for optical stimulation of cells. Sci. Adv. 4, eaat0351 (2018).

9. Zeng, Y. et al. Thermally Conductive Reduced Graphene Oxide Thin Films for Extreme Temperature Sensors. Adv Funct Materials 29, 1901388 (2019).

10. Wang, Y. et al. Reduced graphene oxide film with record-high conductivity and mobility. Materials Today 21, 186–192 (2018).

11. Jaafar, E., Kashif, M., Sahari, S. K. & Ngaini, Z. Study on Morphological, Optical and Electrical Properties of Graphene Oxide (GO) and Reduced Graphene Oxide (rGO). MSF 917, 112–116 (2018).

12. Massicotte, M., Soavi, G., Principi, A. & Tielrooij, K.-J. Hot carriers in graphene – fundamentals and applications. Nanoscale 13, 8376–8411 (2021).

13. Rubio, N. et al. Effect of graphene flake size on functionalisation: quantifying reaction extent and imaging locus with single Pt atom tags. Chem. Sci. 12, 14907–14919 (2021).

14. Kudus, M. H. A., Zakaria, M. R., Akil, H. Md., Ullah, F. & Javed, F. Oxidation of graphene via a simplified Hummers’ method for graphene-diamine colloid production. Journal of King Saud University - Science 32, 910–913 (2020).

15. Al-Dhahebi, A. M., Gopinath, S. C. B. & Saheed, M. S. M. Graphene impregnated electrospun nanofiber sensing materials: a comprehensive overview on bridging laboratory set-up to industry. Nano Convergence 7, 27 (2020).

16. Huang, Y.-L. et al. Self-assembly of graphene onto electrospun polyamide 66 nanofibers as transparent conductive thin films. Nanotechnology 22, 475603 (2011).

17. Engler, A. J., Sen, S., Sweeney, H. L. & Discher, D. E. Matrix Elasticity Directs Stem Cell Lineage Specification. Cell 126, 677–689 (2006).

18. Galie, P. A., Khalid, N., Carnahan, K. E., Westfall, M. V. & Stegemann, J. P. Substrate stiffness affects sarcomere and costamere structure and electrophysiological function of isolated adult cardiomyocytes. Cardiovascular Pathology 22, 219–227 (2013).

19. Jiang, F. X., Yurke, B., Schloss, R. S., Firestein, B. L. & Langrana, N. A. Effect of dynamic stiffness of the substrates on neurite outgrowth by using a DNA-crosslinked hydrogel. Tissue Eng Part A 16, 1873–1889 (2010).

20. Tuft, B. W. et al. Material Stiffness Effects on Neurite Alignment to Photopolymerized Micropatterns. Biomacromolecules 15, 3717–3727 (2014).

21. Li, N. et al. The promotion of neurite sprouting and outgrowth of mouse hippocampal cells in culture by graphene substrates. Biomaterials 32, 9374–9382 (2011).

22. Körner, A., Mosqueira, M., Hecker, M. & Ullrich, N. D. Substrate Stiffness Influences Structural and Functional Remodeling in Induced Pluripotent Stem Cell-Derived Cardiomyocytes. Front. Physiol. 12, (2021).

23. Paşca, A. M. et al. Functional cortical neurons and astrocytes from human pluripotent stem cells in 3D culture. Nat Methods 12, 671–678 (2015).

24. Quadrato, G. et al. Cell diversity and network dynamics in photosensitive human brain organoids. Nature 545, 48–53 (2017).

25. Lian, X. et al. Directed cardiomyocyte differentiation from human pluripotent stem cells by modulating Wnt/β-catenin signaling under fully defined conditions. Nat Protoc 8, 162–175 (2013).

26. Papes, F. et al. Transcription Factor 4 loss-of-function is associated with deficits in progenitor proliferation and cortical neuron content. Nat Commun 13, 2387 (2022).

27. Moore, A. R., Zhou, W.-L., Jakovcevski, I., Zecevic, N. & Antic, S. D. Spontaneous Electrical Activity in the Human Fetal Cortex In Vitro. J Neurosci 31, 2391–2398 (2011).

28. Hogan, M. K., Hamilton, G. F. & Horner, P. J. Neural Stimulation and Molecular Mechanisms of Plasticity and Regeneration: A Review. Front. Cell. Neurosci. 14, (2020).

29. Gleeson, J. G., Lin, P. T., Flanagan, L. A. & Walsh, C. A. Doublecortin Is a Microtubule-Associated Protein and Is Expressed Widely by Migrating Neurons. Neuron 23, 257–271 (1999).

30. Francis, F. et al. Doublecortin Is a Developmentally Regulated, Microtubule-Associated Protein Expressed in Migrating and Differentiating Neurons. Neuron 23, 247–256 (1999).

31. Bifari, F. et al. Complete neural stem cell (NSC) neuronal differentiation requires a branched chain amino acids-induced persistent metabolic shift towards energy metabolism. Pharmacol Res 158, 104863 (2020).

32. Zheng, Y. & Chen, S. Transcriptional precision in photoreceptor development and diseases – Lessons from 25 years of CRX research. Front Cell Neurosci 18, 1347436 (2024).

33. Lakowski, J. et al. Isolation of Human Photoreceptor Precursors via a Cell Surface Marker Panel from Stem Cell-Derived Retinal Organoids and Fetal Retinae. Stem Cells 36, 709–722 (2018).

34. Afanasyeva, T. A. V. et al. A look into retinal organoids: methods, analytical techniques, and applications. Cell Mol Life Sci 78, 6505–6532 (2021).

35. Fligor, C. M. et al. Three-Dimensional Retinal Organoids Facilitate the Investigation of Retinal Ganglion Cell Development, Organization and Neurite Outgrowth from Human Pluripotent Stem Cells. Sci Rep 8, 14520 (2018).

36. Esclapez, M., Tillakaratne, N., Kaufman, D., Tobin, A. & Houser, C. Comparative localization of two forms of glutamic acid decarboxylase and their mRNAs in rat brain supports the concept of functional differences between the forms. J. Neurosci. 14, 1834–1855 (1994).

37. Sadeghianmaryan, A. et al. Electrospinning of Scaffolds from the Polycaprolactone/Polyurethane Composite with Graphene Oxide for Skin Tissue Engineering. Appl Biochem Biotechnol 191, 567– 578 (2020).

38. Chen, X. et al. Characteristics and toxicity assessment of electrospun gelatin/PCL nanofibrous scaffold loaded with graphene in vitro and in vivo. Int J Nanomedicine 14, 3669–3678 (2019).

39. Ghasemi, A. et al. Studying the Potential Application of Electrospun Polyethylene Terephthalate/Graphene Oxide Nanofibers as Electroconductive Cardiac Patch. Macromolecular Materials and Engineering 304, 1900187 (2019).

40. Ruan, K., et al. Improved thermal conductivities in polystyrene nanocomposites by incorporating thermal reduced graphene oxide *via* electrospinning-hot press technique. Composites Communications 10, 68–72 (2018).

41. Oh, S. H. & Lee, J. H. Hydrophilization of synthetic biodegradable polymer scaffolds for improved cell/tissue compatibility. Biomed Mater 8, 014101 (2013).

42. Domingos, M. et al. Improved osteoblast cell affinity on plasma-modified 3-D extruded PCL scaffolds. Acta Biomaterialia 9, 5997–6005 (2013).

43. Ding, J. et al. Electrospun polymer biomaterials. Progress in Polymer Science 90, 1–34 (2019).

44. Chen, L., Yan, C. & Zheng, Z. Functional polymer surfaces for controlling cell behaviors. Materials Today 21, 38–59 (2018).

45. Zhao, G. et al. Reduced graphene oxide functionalized nanofibrous silk fibroin matrices for engineering excitable tissues. NPG Asia Mater 10, 982–994 (2018).

46. Metwally, S. & Stachewicz, U. Surface potential and charges impact on cell responses on biomaterials interfaces for medical applications. Materials Science and Engineering: C 104, 109883 (2019).

47. Safaeijavan, R., Soleimani, M., Divsalar, A., Eidi, A. & Ardeshirylajimi, A. Comparison of random and aligned PCL nanofibrous electrospun scaffolds on cardiomyocyte differentiation of human adipose-derived stem cells. Iran J Basic Med Sci 17, 903–911 (2014).

48. Xie, J. et al. ‘Aligned-to-random’ nanofiber scaffolds for mimicking the structure of the tendon-to-bone insertion site. Nanoscale 2, 923–926 (2010).

49. Yu, C.-C. et al. Random and aligned electrospun PLGA nanofibers embedded in microfluidic chips for cancer cell isolation and integration with air foam technology for cell release. Journal of Nanobiotechnology 17, 31 (2019).

50. Xie, J., MacEwan, M. R., Li, X., Sakiyama-Elbert, S. E. & Xia, Y. Neurite Outgrowth on Nanofiber Scaffolds with Different Orders, Structures, and Surface Properties. ACS Nano 3, 1151– 1159 (2009).

51. Yang, F., Murugan, R., Wang, S. & Ramakrishna, S. Electrospinning of nano/micro scale poly(l-lactic acid) aligned fibers and their potential in neural tissue engineering. Biomaterials 26, 2603–2610 (2005).

52. Soles, A. et al. Extracellular Matrix Regulation in Physiology and in Brain Disease. Int J Mol Sci 24, 7049 (2023).

53. Sood, D. et al. Fetal brain extracellular matrix boosts neuronal network formation in 3D bioengineered model of cortical brain tissue. ACS Biomater Sci Eng 2, 131–140 (2016).

54. Sernagor, E. Retinal Development: Second Sight Comes First. Current Biology 15, R556–R559 (2005).

55. Zeck, G., Lambacher, A. & Fromherz, P. Axonal transmission in the retina introduces a small dispersion of relative timing in the ganglion cell population response. PLoS One 6, e20810 (2011).

56. Bose, M. et al. Effect of the environment on the dendritic morphology of the rat auditory cortex. Synapse 64, 97–110 (2010).

57. Greifzu, F. et al. Environmental enrichment extends ocular dominance plasticity into adulthood and protects from stroke-induced impairments of plasticity. Proceedings of the National Academy of Sciences 111, 1150–1155 (2014).

58. Engineer, N. D. et al. Environmental enrichment improves response strength, threshold, selectivity, and latency of auditory cortex neurons. J Neurophysiol 92, 73–82 (2004).

59. Feldman, D. E. & Brecht, M. Map plasticity in somatosensory cortex. Science 310, 810–815 (2005).

60. Bertolesi, G. E., Hehr, C. L. & McFarlane, S. Wiring the retinal circuits activated by light during early development. Neural Dev 9, 3 (2014).

61. Bonezzi, P. J., Tarchick, M. J., Moore, B. D. & Renna, J. M. Light drives the developmental progression of outer retinal function. J Gen Physiol 155, e202213262 (2023).

62. Zhang, J., Cao, H.-Y., Wang, J.-Q., Wu, G.-D. & Wang, L. Graphene Oxide and Reduced Graphene Oxide Exhibit Cardiotoxicity Through the Regulation of Lipid Peroxidation, Oxidative Stress, and Mitochondrial Dysfunction. Front. Cell Dev. Biol. 9, 616888 (2021).

63. Gurunathan, S., Woong Han, J., Abdal Daye, A., Eppakayala, V. & Kim, J. Oxidative stress-mediated antibacterial activity of graphene oxide and reduced graphene oxide in Pseudomonas aeruginosa. IJN 5901 (2012) doi:10.2147/IJN.S37397.

64. Kang, Y. et al. Oxidation of Reduced Graphene Oxide *via* Cellular Redox Signaling Modulates Actin-Mediated Neurotransmission. ACS Nano 14, 3059–3074 (2020).

65. Sharow, K. A., Temkin, B. & Asson-Batres, M. A. Retinoic acid stability in stem cell cultures. The International Journal of Developmental Biology 56, 273–278 (2012).

66. Czuba, L. C., Zhong, G., Yabut, K. & Isoherranen, N. Analysis of Vitamin A and Retinoids in Biological Matrices. Methods Enzymol 637, 309–340 (2020).

67. Fu, P. P. et al. Photodecomposition of Vitamin A and Photobiological Implications for the Skin†. Photochemistry and Photobiology 83, 409–424 (2007).

68. Kelley, M. W., Turner, J. K. & Reh, T. A. Retinoic acid promotes differentiation of photoreceptors in vitro. Development 120, 2091–2102 (1994).

69. Sanjurjo-Soriano, C. et al. Retinoic acid delays initial photoreceptor differentiation and results in a highly structured mature retinal organoid. Stem Cell Res Ther 13, 478 (2022).

70. Li, X., Zhang, L., Tang, F. & Wei, X. Retinal Organoids: Cultivation, Differentiation, and Transplantation. Front. Cell. Neurosci. 15, (2021).

71. Zhong, X. et al. Generation of three-dimensional retinal tissue with functional photoreceptors from human iPSCs. Nat Commun 5, 4047 (2014).

72. Fernando, M. et al. Differentiation of brain and retinal organoids from confluent cultures of pluripotent stem cells connected by nerve-like axonal projections of optic origin. Stem Cell Reports 17, 1476–1492 (2022).

73. Zhang, Z., Liu, Y., Lin, S. & Wang, Q. Preparation and properties of glutaraldehyde crosslinked poly(vinyl alcohol) membrane with gradient structure. J Polym Res 27, 228 (2020).

74. Kaushik, G., Fuhrmann, A., Cammarato, A. & Engler, A. J. In Situ Mechanical Analysis of Myofibrillar Perturbation and Aging on Soft, Bilayered Drosophila Myocardium. Biophysical Journal 101, 2629–2637 (2011).

75. Fitzgerald, M. Q. et al. Generation of ‘semi-guided’ cortical organoids with complex neural oscillations. Nat Protoc 1–27 (2024) doi:10.1038/s41596-024-00994-0.

76. Andrews, Simon. FastQC: A Quality Control tool for High Throughput Sequence Data.

77. 77. Krueger, F. Trim Galore! https://www.bioinformatics.babraham.ac.uk/projects/trim_galore/.

78. Bray, N. L., Pimentel, H., Melsted, P. & Pachter, L. Near-optimal probabilistic RNA-seq quantification. Nat Biotechnol 34, 525–527 (2016).

79. Soneson, C., Love, M. I. & Robinson, M. D. Differential analyses for RNA-seq: transcript-level estimates improve gene-level inferences. Preprint at 10.12688/f1000research.7563.1 (2016).

80. Leek, J. T., Johnson, W. E., Parker, H. S., Jaffe, A. E. & Storey, J. D. The sva package for removing batch effects and other unwanted variation in high-throughput experiments. Bioinformatics 28, 882–883 (2012).

81. Kolde, R. pheatmap: Pretty Heatmaps. (2019).

82. Ritchie, M. E. et al. limma powers differential expression analyses for RNA-sequencing and microarray studies. Nucleic Acids Res 43, e47 (2015).

83. Law, C. W., Chen, Y., Shi, W. & Smyth, G. K. voom: precision weights unlock linear model analysis tools for RNA-seq read counts. Genome Biology 15, R29 (2014).

84. Gao, C.-H., Yu, G. & Cai, P. ggVennDiagram: An Intuitive, Easy-to-Use, and Highly Customizable R Package to Generate Venn Diagram. Front. Genet. 12, (2021).

85. Gao, C.-H., et al. ggVennDiagram: Intuitive Venn diagram software extended. iMeta 3, e177 (2024).

86. Wickham, H. Ggplot2: Elegant Graphics for Data Analysis. (2016).

87. Wu, T. et al. clusterProfiler 4.0: A universal enrichment tool for interpreting omics data. Innovation 2, (2021).

88. Yu, G., Wang, L.-G., Han, Y. & He, Q.-Y. clusterProfiler: an R Package for Comparing Biological Themes Among Gene Clusters. OMICS: A Journal of Integrative Biology 16, 284–287 (2012).

89. Larmarange, J. ggstats: Extension to ggplot2 for Plotting Stats. https://larmarange.github.io/ggstats/ (2024).

90. Kassambara, A. ggpubr: ‘ggplot2’ Based Publication Ready Plots. (2023).

